# Yeast galactokinase in closed conformation can switch between catalytic and signal transducer states

**DOI:** 10.1101/2022.03.07.483271

**Authors:** Nandinee Giri, Paike Jayadeva Bhat

## Abstract

*S.cerevisiae* galactokinase (*Sc*Gal1p), in closed conformation catalyzes the phosphorylation of galactose to galactose 1-phopshate using ATP as the phosphate donor as well as allosterically activates the *GAL* genetic switch in response to galactose and ATP as ligands. How both kinase and signaling activities of *Sc*Gal1p are associated with closed conformation of the protein is not understood. Conformational sampling of *Sc*Gal1p indicated that this protein samples closed kinase and closed non-kinase conformers. Closed non-kinase conformers are catalytically incompetent to phosphorylate galactose and act as a bonafide signal transducer. It was observed that toggling of side chain of highly conserved K266 of *Sc*Gal1p between S171and catalytic base D217 is responsible for transitioning of *Sc*Gal1p between signal transducer and kinase states. Interestingly in *Sc*Gal3p, the paralog of *Sc*Gal1p, which has only signal transduction activity and lacks kinase activity, a H bond between a non-conserved Y433, and a highly conserved Y57, gets broken during MD simulation. The corresponding H-bond present in *Sc*Gal1p between residues Y441 and Y63 respectively, remains intact throughout the simulations of *Sc*Gal1p.Therefore, we predicted that K266 and Y441 have a role in bifunctionality of *Sc*Gal1p. To test the above predictions, we monitored the signaling and kinase activity of *Sc*Gal1_K266R_p and *Sc*Gal1_Y441A_p variants. Signaling activity increased in both *Sc*Gal1_Y441A_p and *Sc*Gal1_K266R_p variants as compared to *Sc*Gal1_wt_p, whereas the kinase activity increased in *Sc*Gal1_Y441A_p, but decreased in *Sc*Gal1_K266R_p Based on the above, we propose that K266 and Y441 are crucial for conferring bifunctionality to *Sc*Gal1p.

**Author summary:** Galactokinase of *S.cerevisiae*(*Sc*Gal1p), the first enzyme of Leloir pathway of galactose metabolism, phosphorylates galactose using ATP as the phosphate donor. *Sc*Gal1p also functions as a signal transducer of *GAL* regulon wherein galactose and ATP allosterically activate galactokinase. The active form of galactokinase, then sequesters the repressor *Sc*Gal80p, to activate the *GAL* switch. *Sc*Gal1p has a single site each for binding to galactose and ATP. How *Sc*Gal1p, a monomeric protein, performs the above two mutually exclusive activities using the same set of substrates/ligands, with the same site acting as the active site for enzymatic activity as well as allosteric site for signal transduction activity is unclear. Our findings are that this protein has a distinct conformational state for functioning as a signal transducer and a distinct conformational state for functioning as a kinase. A highly conserved lysine residue (K266) present only in fungal galactokinases, triggers the interconversion between catalysis and signal transduction states. This interconversion is subdued by H bond between Y441 and Y63. These studies suggest that the two activities of ScGal1p are fine tuned by evolution to regulate metabolism through transcriptional control.

## Introduction

The text book concept of molecular biology, that one sequence-one protein-one function [1, 2, 3] has now been largely replaced by one protein more functions [4, 5]. Mechanisms such as gene fusion /domain fusion events [6], alternative splicing [7], programmed frame shifting [8] and phenotypic mutations [9, 10] introduce multifunctionality into a protein, resulting in functional diversification. Additionally, it was recognized that a polypeptide can code for two or more than two, often disparate biochemical activities, a phenomenon originally described as gene sharing [11], but now commonly referred to as moonlighting [12]. In moonlighting proteins, autonomous biochemical activities emanate from a single structure and therefore not to be confused with multifunctionality discussed above. That is, moonlighting proteins do not have autonomous structural domains to carryout distinct moonlighting activities. And yet, to be qualified as a moonlighting protein, different activities of moonlighting proteins should be mutationally separable [13], forcing us to re-evaluate our current understanding of structure-function paradigm of proteins. Moonlighting proteins have attracted considerable attention because of their ubiquity [14], involvement in diverse cellular processes [15] role in protein evolution following gene duplication [16, 17] and have also been implicated in many diseases processes [18, 19].

In general, moonlighting activities have no apparent functional link with the canonical/primary activity. In fact, a large number of metabolic enzymes involved in central carbon and nitrogen metabolism are now known to be moonlighting [20]. For example, Phosphoglucoseisomerase, the first moonlighting protein to be identified, also functions as a neurotropic factor [21]. Mutation that abolishes the enzyme activity causes haemolytic anaemia while mutations that abolish both the activities cause haemolytic anaemia with neurological defects [22]. Aconitase of yeast, an Fe-S cluster containing enzyme of citric acid cycle, participates in the maintenance of mtDNA, an activity independent of its enzymatic activity [23].Thus, identification, functional annotation, understanding the evolutionary origin and above all deciphering the underlying molecular mechanism of moonlighting activities is fraught with problems.

A more intriguing case of moonlighting is exhibited by galactokinase of *S.cerevisiae*, where in, galactose and ATP serve as substrates for its enzyme activity while the same set of molecules allosterically activate galactokinase to function as a signal transducer of the *GAL* genetic switch [24,25,26]. Thus, the fundamental question as to how yeast galactokinase, a monomeric protein, carries out two mutually exclusive biochemical activities in response to the same set of substrates/ligands remains unanswered.

*S.cerevisiae GAL*1 (*ScGAL1*) encoded galactokinase (*Sc*Gal1p),the first enzyme of Leloir pathway of galactose metabolism, is one of the earliest proteins to be identified as moonlighting [27, 28, 29]. The canonical biochemical activity of *Sc*Gal1p is to catalyse the phosphorylation of galactose using ATP as the phosphate donor, resulting in the formation of galactose 1-phosphate [30]. The function of this activity is to feed galactose into the glycolytic pathway, which is used for generating metabolic energy and intermediates. The second biochemical activity is to sequester *Sc*Gal80p, the repressor of the *GAL* switch, in response to galactose and ATP [26, 31], which serve as ligands for allosteric activation of *Sc*Gal1p. The function of this biochemical activity is to alleviate the *Sc*Gal80p mediated transcriptional repression, thereby allowing *Sc*Gal4p to activate the transcription of galactose metabolizing genes [27, 28, 29].

It needs to be noted that the bonafide signal transducer of the *GAL* genetic switch of *S.cerevisiae* is *Sc*Gal3p. Upon addition of galactose,the signal transducer protein (*Sc*Gal3p) binds to galactose and ATP, thereby attaining the active state. The active state of the signal transducer protein binds to *Sc*Gal80p, relieving its repression on *Sc*Gal4p, thereby allowing the expression of proteins responsible for galactose metabolism [24, 25, 26]. *ScGAL1* and *ScGAL3* arose by duplication from an ancestral bifunctional galactokinase [32, 33]. Consequently,*ScGAL3* encoded protein*Sc*Gal3p,shares 74% identity and 90% similarity with the amino acid sequence of *Sc*Gal1p [25]. *Sc*Gal3p evolved into the bonafide signal transducer of the *GAL* regulon wherein it lost its enzymatic activity completely. On the other hand *Sc*Gal1p was able to retain both its enzymatic as well as signal transduction activity. Thus, in a wild type *S.cerevisaie* strain, once the transcriptional induction of *GAL* genes is ensued by galactose, the induced state can be sustained even after the withdrawal of *Sc*Gal3p function, as long as galactose is available [34, 35]. In a wild type strain, once the induction occurs, the maintenance of the induced state is sustained by *Sc*Gal1p but not *Sc*Gal3p [36].

Structure of *Sc*Gal1p in presence of galactose and non-hydrolysable analogue of ATP, has been solved at a resolution of 3Å [37]. Structure of *Sc*Gal3p both in open and closed conformation has been solved at a resolution of 1.5Å [38]. In the absence of galactose and ATP, *Sc*Gal3p attains the open conformation. In open conformation or the inactive conformation, *Sc*Gal3p cannot bind to *Sc*Gal80p and hence cannot function as a signal transducer. Upon binding to galactose and ATP, *Sc*Gal3p attains the closed or active conformation, which in turns binds to *Sc*Gal80p allowing *Sc*Gal3p to function as a signal transducer [38]. Above studies indicated that both proteins exhibit a bilobal structure with galactose and ATP binding in between the N and C domains. It was proposed that the absence of kinase activity in *Sc*Gal3p is due to the orientation of the anomeric -OH of galactose in a configuration not compatible with catalysis [38]. Absence of kinase activity in *Sc*Gal3p is also attributed to a deletion of SA dipeptide, corresponding to S171 and A172 of *Sc*Gal1p [39].

Over the past three decades, several independent studies have introduced substitutions in *ScGAL1* [29, 40], *ScGAL3* [41] and *KlGAL1* (*K. lactis GAL1*, an orthologue of *ScGAL1* in *Kluyveromyces lactis* is also a moonlighting protein and has been studied extensively) [28, 42] to understand the mechanistic basis of their biochemical activities (see Table 1 for details). However, these studies do not shed light on how galactose and ATP serve as substrates for catalysis as well as ligands for allosteric activation of *Sc*Gal1p. Further, it needs to be pointed out that from the crystal structure of *Sc*Gal1p it was observed that this protein has a single site each for binding to galactose and ATP. That is, the allosteric and orthosteric sites are the same. Hence the bifunctionality observed in *Sc*Gal1p becomes all the more intriguing as it performs two functions using ATP and galactose as substrates for enzymatic activity as well as ligands for the signal transduction activity with the allosteric site for regulatory function completely overlapping with orthosteric site for catalytic activity.

**Table 1:**
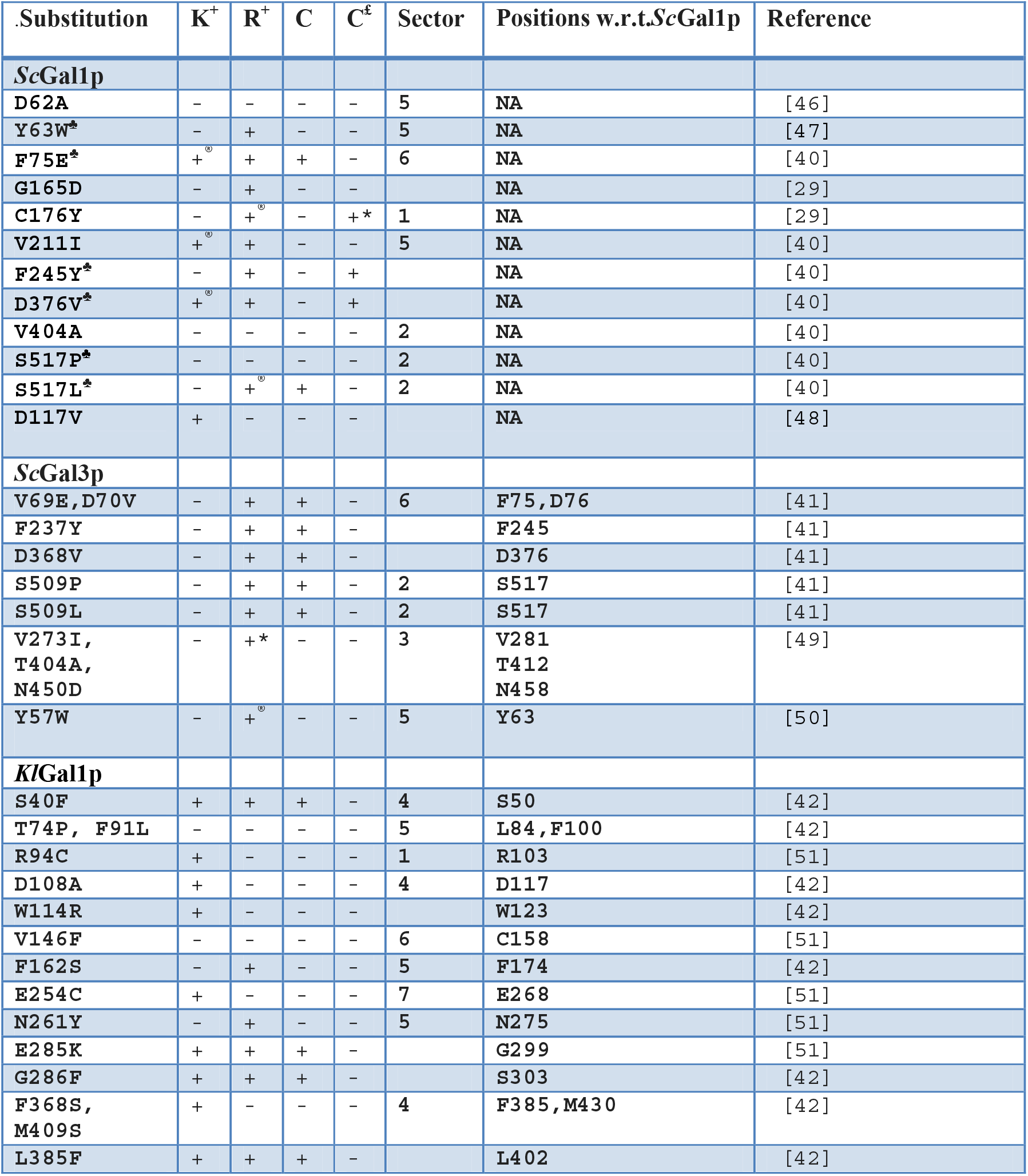
List of substitutions in KlGal1p, ScGal1p and ScGal3p. K**^+^** and R**^+^** indicate that the mutant has retained the kinase and regulatory activities respectively. ♣indicates constitutive mutations initially identified in ScGal3p were introduced into ScGal1p. C indicates the constitutive mutants that are responsive to galactose. C^£^ indicates the constitutive mutant variants that exhibit galactose independent induction activity. *indicates mutant of ScGal3p/ScGal1p that functions as a signal transducer only upon over-expression. ® indicates reduction in the activity as compared to wild type. NA indicates not applicable.

We carried out computational studies to hone into the possible amino acid residues that might participate in the transition between catalysis and allosteric activation. From the above analysis, we identified K266 and Y441 for mutational analysis. Based on phenotypic analysis, growth kinetics and biochemical studies of *Sc*Gal1_wt_p, *Sc*Gal1_K266R_p and *Sc*Gal1_Y441A_p variants in a strain otherwise deleted for *GAL3* (*gal3Δ* background), we propose that K266 acts as a pivot or as a ‘swinging arm’ to tip the balance between catalytic and signalling conformations. On the other hand, a H bond between Y441, an evolutionarily non-conserved residue and Y63, an evolutionarily conserved residue, dampens this transition. Based on our results and previous reports, we discuss how evolutionary forces tinker with molecules [43] to fine tune the cellular response to the vagaries of environmental fluctuations.

## Results

### Sector analysis of galactokinase of *S. cerevisiae*

Studies carried out over the past three decades have identified 30 amino acid substitutions which alter kinase, signalling or both activities in *Sc*Gal1p, *Kl*Gal1p and *Sc*Gal3p (see Table 1 for details). Inspection of the crystal structure in light of the above substitutions, do not provide clues to understand the mechanistic basis of bifunctionality. We surmised that bifunctionality in all likelihood is a consequence of residue-residue interaction leading to altered protein dynamics. Such interacting residues present at distant locations in the three dimensional structures generally co-evolve. Statistical coupling analysis (SCA), has been successfully applied to identify functionally relevant residues as sectors of co-evolving residues [44].

We carried out SCA of yeast galactokinase to identify sectors, if any, involved in bifunctionality. We detected 7 sectors in *Sc*Gal1p (see Figure S1). Of the 29 experimentally identified residues, only 18 could be mapped to different sectors. Sectors 4, 5 and 6 accounts for 65% of experimentally identified residues as important for function (see Table S1). It turns out that of these 29 experimentally identified substitutions, 5 of them confer regulatory positive, kinase negative(R^+^K^-^) phenotype either to *Sc*Gal1p or *Kl*Gal1p.That is, the signal transduction function remains similar to that wild type *Sc*Gal1p/*Kl*Gal1p as a consequence of these substitutions. Of these 5 residues, 4 belong to sector 5 and 1 does not belong to any sector.Sector 5 also consists of two other experimentally identified substitutions, which result in inactivation of both the functions of either *Sc*Gal1p or *Kl*Gal1p

Inspection of the crystal structure of *Sc*Gal1p indicated that residues present in sector 5 participate in galactose binding and form the first shell of polar interaction [37] with the binding residues. Given that mutants that are part of this sector result in abrogation of kinase activity in both *Sc*Gal1p and *Kl*Gal1p, with their signal transduction activity remaining intact, we inferred that sector 5 has a role in kinase function. This inference gets further strengthened by virtue of the fact that although sector 5 accommodates both galactose and ATP, the majority of experimentally identified substitutions residing in sector 5 impair only the phosphotransfer reaction (That is, they are able to function as signal transducer in response to galactose). This implies that these residues do not inhibit binding of galactose and ATP to *Sc*Gal1p or *Kl*Gal1p.

Residue K266, which was previously shown to re-orient during MD simulation run of 50 ns [45], also resides in sector 5.The dipeptide _171_SA_172_ that confers kinase activity to *Sc*Gal3p [39], resides in sector 6. Interestingly, the crystal structure of *Sc*Gal1p shows that K266 forms H bond interaction with S171 (the corresponding residue in *Sc*Gal3p is absent). Above observations suggested that sector 5 and sector 6 cross talk through the interaction of K266 and S171 (figure1). While the role of other sectors is not clear, it is possible that they could participate in functions such as stability, allostery. Nevertheless, the results obtained from SCA hinted that the residues of Sector 5 and 6 could have a role in bifunctionality.

### Preliminary MD simulation results indicated that the closed structure of *Sc*Gal1p can exist in kinase and non-kinase state

To probe the interaction between K266 and S171, we carried out MD simulation for a period of 10ns. The starting structure for MD simulation is the crystal structure of *Sc*Gal1p [37] in which AMPPNP conformation is compact, a conformation not suitable for catalysis, because the distance between terminal phosphate of AMPPNP and anomeric -OH group of galactose is more than 6Å. In this conformation, K266 is in H bond interaction with S171 but not with D217 (Figure 2A), which is thought to act as a catalytic base [38]. For carrying out molecular dynamic simulation of *Sc*Gal1p, we replaced AMPPNP with ATP. As the simulation progressed, the side chain of K266 started orienting towards D217. That is, H-bond interaction occurs with K266 and D217 concomitant with the change in ATP conformation from compact to extended state. If the distance between terminal phosphate and the adenine ring of ATP is less than 9.0 Å, ATP is said to exist in compact conformation, if more it is said to be in extended conformation [52] (see figure 2A and 2C). This transition of ATP conformation results in a decrease in the distance between terminal phosphate of ATP and anomeric -OH of galactose to as low as 4.7Å (Figure 2C). During the above simulation, the lip distance (i.e. the distance between D106 andV383) remained less than 35Å, suggesting that the reorientation of K266 does not increase the lip distance beyond 35Å, implying that the structure is still in closed conformation [38](Figure 2 B and D). This suggested that movement of the side chain of K266 converts the non-catalytic state to the catalytic state without causing an increase in the lip distance. If so, the structure before the movement of K266 towards D217 could as well represent the signal transduction state of *Sc*Gal1p. That is, under these conditions the terminal phosphate of ATP is not within a distance compatible for catalysis (Figure 2A & C).

**Figure 1:**
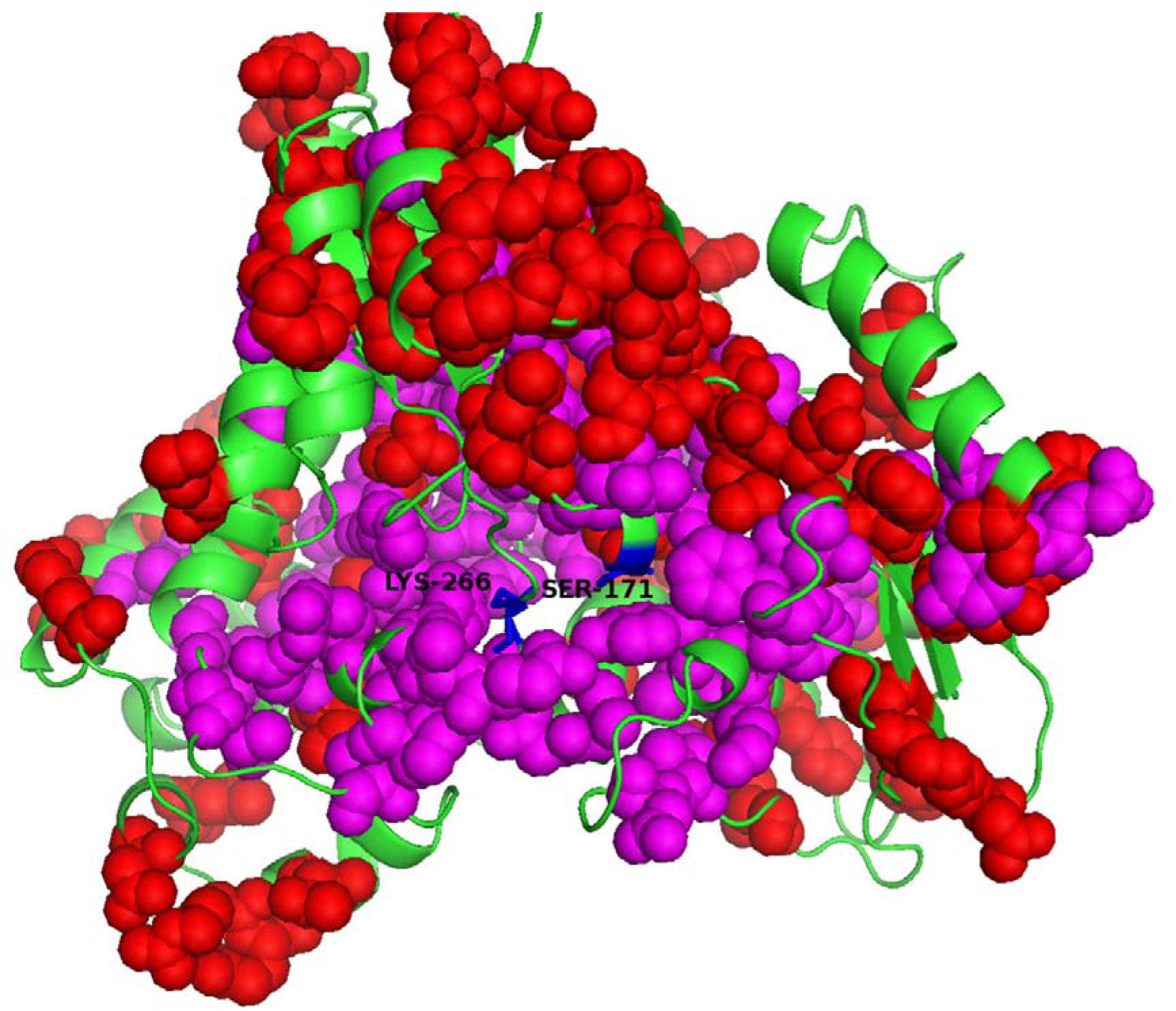
Mapping of residues of sector 5 and 6 onto *Sc*Gal1p structure. Residues of sector 5 are coloured magenta and sector 6 are coloured red. K266 (part of sector 5) and S171(part of sector 6) are coloured in blue. K266 and S171 interact via a H-bond [37].

**Figure 2:**
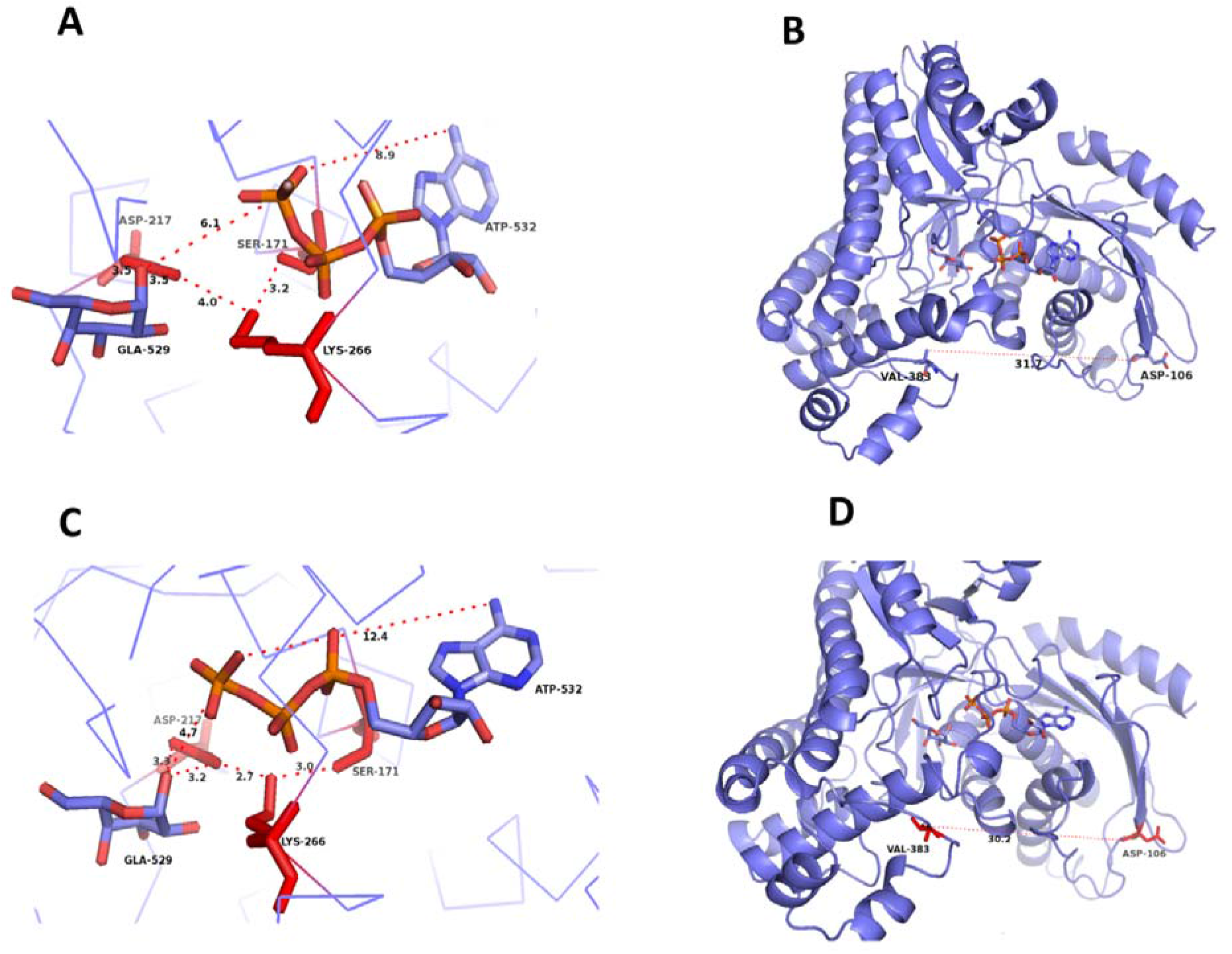
A snapshot of the orientation of various residues in *Sc*Gal1p. Figures A and C show the structural configurations of S171, K266, D217, ATP and galactose in non kinase and kinase conformations of *Sc*Gal1p respectively. K266,S171, D217 are coloured red whereas ATP and galactose are coloured purple. Figures B and D indicate that the corresponding *Sc*Gal1p structures remain in closed conformation as indicated by the lip distance. The dashed lines represent the indicated distances in Å

Thus, we surmise that *Sc*Gal1p exists at least in three conformational states: catalytic state, signal transducer state (both of which are in closed conformation) and open state which is neither catalytic nor signal transducer state. The existence of open state is rationalised based on the observation that overexpression of *Sc*Gal1p leads to constitutive expression [29] and in analogy with the open conformation of *Sc*Gal3p [38, 49]. Thus, as a starting point, the orientation of the side chain of K266 either towards or away from D217 was considered as the discriminating feature between the kinase and signal transduction respectively.

### S171 of SA dipeptide keeps K266 in vicinity of D217 which is required for the catalytic activity of *Sc*Gal1p

From the 10ns simulation, the role of S171 in kinase activity could not be deciphered. To further probe whether or not K266 is important for transition of *Sc*Gal1p to kinase state and to decipher the role of S171, we carried out simulation of *Sc*Gal1p, *Sc*Gal3p and*Sc*Gal3-SAp for 150ns. The starting structure of *Sc*Gal3-SAp was obtained by inserting SA dipeptide in the crystal structure of *Sc*Gal3p. Here,*Sc*Gal3p and *Sc*Gal3-SAp served as negative and positive control respectively. This is because,*Sc*Gal3p lacks kinase activity but retains signalling activity while *Sc*Gal3-SAp has signaling activity and weak kinase activity. During this simulation, we monitored the distance between the following residues: (i) distance between terminal phosphate of ATP and anomeric -OH group of galactose (ii) distance between D217 and anomeric -OH group of galactose (iii) distance of ε amino group of K266 from D217 and S171. Please note that the numbering of residues mentioned here is with respect to *Sc*Gal1p. The corresponding residues in *Sc*Gal3p and *Sc*Gal3-SAp are shown in figure 3C.

**Figure 3:**
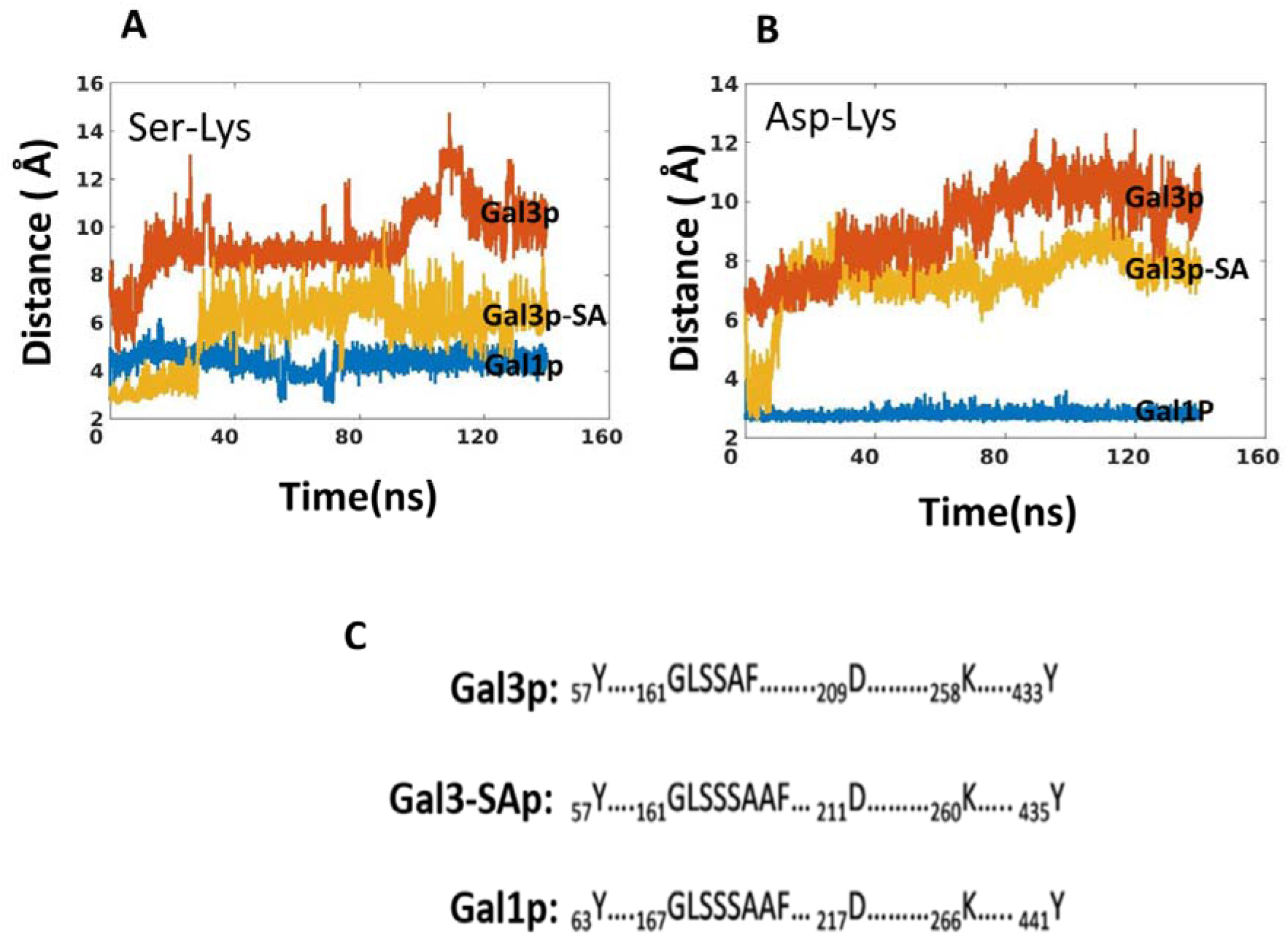
Trajectories of lysine residue in *Sc*Gal1p, *Sc*Gal3p and *Sc*Gal3p-SAp. Panel A shows the variation of distance between serine and lysine residue in *Sc*Gal3p, *Sc*Galp and *Sc*Gal3-SAp. The numbering of serine is S164, S165 and S171 in *Sc*Gal3p, *Sc*Gal3-SAp and *Sc*Gal1p respectively (see figure 3,panel C and text for more). Panel B shows the variation of distance between catalytic aspartate and lysine residue in *Sc*Gal3p, *Sc*Galp and *Sc*Gal3-SAp. The numbering of lysine residue is K258, K260 and K266 in *Sc*Gal3p, *Sc*Gal3-SAp and *Sc*Gal1p respectively(see figure 3, panel C). The numbering of catalytic aspartate is D209, D211 and D217 in *Sc*Gal3p, *Sc*Gal3p-SAp and *Sc*Gal1p respectively(see figure 3, panel C). details). Panel C shows the numbering system used to refer to the relevant corresponding residues in *Sc*Gal3p and *Sc*Gal3-SAp.

The first two order parameters tell us whether the conformation is compatible with catalysis or not. Third order parameter was used to find out whether K266 plays any role in transition of *Sc*Gal1p between kinase and signalling states. Note that, unlike D217, which is thought to be a catalytic base, no specific role has been attributed to S171. Nevertheless, given its importance in imparting kinase activity to *Sc*Gal3p [39], we thought it is prudent to use S171 as a reference for monitoring the movement of K266 side chain.

During these simulations, it was observed that in *Sc*Gal3p, K258 (corresponding to K266 of *Sc*Gal1p, see Figure 3, Panel C) moves away from S164 (see Figure 3 Panel A and Figure S4, Panel A & B). Since SA dipetide is missing in *Sc*Gal3p, we measured the distance of K258 from S164. The residue corresponding to S164 of *Sc*Gal3p is S170 in *Sc*Gal1p (see Figure 3, Panel C). However, S164 in the crystal structure of *Sc*Gal3p is structurally analogous to S171 (S of SA dipetide) of *Sc*Gal1p. In case of *Sc*Gal3-SAp, the distance between K260 (corresponding to K266 of *Sc*Gal1p, see Figure 3, Panel C) and S165 (corresponding to S171 of *Sc*Gal1p, see Figure 3, Panel C) is lesser as compared to distance between K258 and S164 of *Sc*Gal3p (see Figure 3, Panel A and Figure S2, Panel A & B). Thus, addition of SA dipeptide in *Sc*Gal3p (i.e.*Sc*Gal3-SAp) seems to restrict the motion of K260. In comparison to both *Sc*Gal3p and *Sc*Gal3-SAp, the distance between K266 and S171 in *Sc*Gal1p remains the lowest (see Figure 3, Panel A and Figure S3). Note that, the distance between lysine (i.e. K266 of *Sc*Gal1p, K260 of *Sc*Gal3-SAp and K258 of *Sc*Gal3p) and serine (i.e. S171 of ScGal1p, S165 of ScGal3-SAp and S164 of *Sc*Gal3p) was determined by measuring the distance between NZ atom of lysine and OG atom of serine (names of the atom are as per the PDB format).

From this, we inferred that S171 (i.e. S of SA dipeptide) restricts the motion of K266 in *Sc*Gal1p and keeps it in the vicinity of the active site (i.e. D217 of *Sc*Gal1p). S171 appears to hold K266 in a standby mode for catalysis, thus serving as an anchor. In absence of S171, the proper positioning of K266 can’t occur and it would move further away from active site (D217) of *Sc*Gal1p. Hence, the putative role of S171 could be to govern the motion of K266.

Consequently, it was observed that the distance between K266 and D217 remained lowest in *Sc*Gal1p (see Figure 3, Panel B and Figure S3), followed by *Sc*Gal3-SAp (see Figure 3, Panel B and Figure S2, Panel A & B) and then *Sc*Gal3p (see Figure 3, Panel B and Figure S4, Panel A & B). In *Sc*Gal3-SAp, the residues corresponding to K266 and D217 are K260 and D211 respectively, whereas in ScGal3p, the corresponding residues are K258 and D209 respectively (see Figure 3, Panel C). In case of *Sc*Gal1p, the distance between K266 and D217 always remained within H-bond range (see Figure 3, Panel B). The distance between lysine (i.e. K266 of ScGal1p, K260 of *Sc*Gal3-SAp and K258 of *Sc*Gal3p) and catalytic aspartate (i.e. D217 of *Sc*Gal1p, D211 of *Sc*Gal3-SAp and D209 of *Sc*Gal3p) was measured by measuring the distance between NZ atom of lysine and OD1 and OD2 atom of aspartate. The minimum of these two distances was taken as the distance between afore-mentioned lysine residue and catalytic aspartate residue (names of the atom are as per the PDB format).

The vicinity of K266 to D217 (see Figure 3, Panel B) allows *Sc*Gal1p to remain in the kinase state as the distance between D217 and anomeric hydroxyl group of galactose in *Sc*Gal1p always remained within the range of H-bond interaction (see Figure 4, Panel A and Figure S3). In contrary to this in *Sc*Gal3p, H-bond never formed between D209 and anomeric hydroxyl group of galactose (see Figure 4, Panel A) as K258 was distant from D209 (see Figure 3 Panel B and Figure S4, Panel A & B). On the other hand, in case of *Sc*Gal3-SAp, as expected, it was observed that H-bond interaction subsequently forms between anomeric hydroxyl group of galactose and D211 (see Figure 4, Panel A and Figure S2, Panel A & B). Recall that the distance between K260 and D211 in *Sc*Gal3-SAp remain lower that what was observed in *Sc*Gal3p, but higher than *Sc*Gal1p (see Figure 3, Panel B). The distance between catalytic aspartate (i.e. D217 of ScGal1p, D211 of *Sc*Gal3-SAp and D209 of *Sc*Gal3p) and anomeric hydroxyl group of galactose was measured by measuring the distance of OD1 and OD2 atom of aspartate from anomeric hydroxyl group of galactose. The minimum of these two distances was taken as the distance between anomeric hydroxyl group of galactose and catalytic aspartate (names of the atom are as per the PDB format).

**Figure 4:**
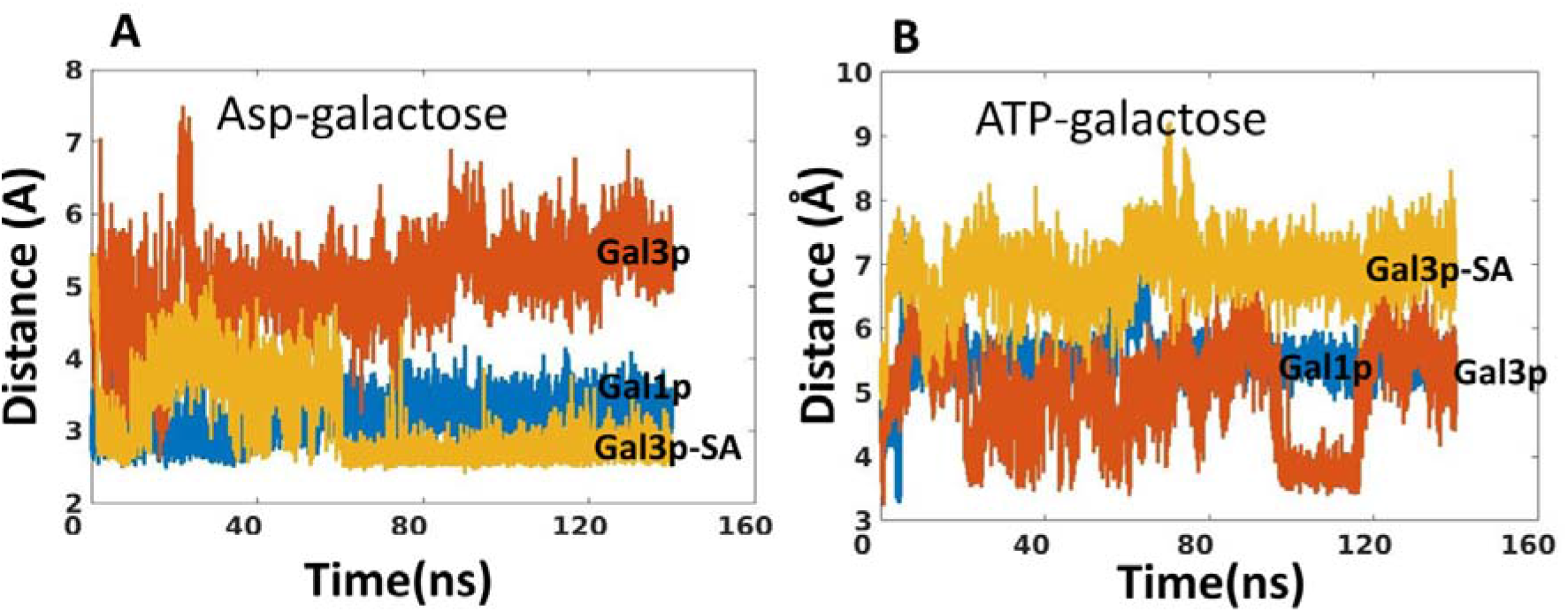
Trajectories of different catalytic entities in *Sc*Gal1p, *Sc*Gal3p and *Sc*Gal3p-SAp. Panel A shows the distance between catalytic aspartate and anomeric -OH group of galactose in *Sc*Galp, *Sc*Gal3p, and *Sc*Gal3p-SAp, as a function of time. The numbering of catalytic aspartate is D209, D211 and D217 in *Sc*Gal3p, *Sc*Gal3p-SAp and *Sc*Gal1p respectively (see Figure 3 Panel C for details). Panel B shows the distance between terminal phosphate of ATP and anomeric -OH group of galactose in *Sc*Gal1p, *Sc*Ga3p and *Sc*Gal3p-SAp, as a function of time.

Based on the above, we hypothesize that S171 restricts the motion of K266 and keeps it within close proximity of D217, thereby allowing *Sc*Gal1p to transition to kinase state. This is because the vicinity of K266 toD217 maintains the geometry of D217 and allows the formation H-bond with anomeric hydroxyl group of galactose and D217.

### H-bond between D217 and anomeric hydroxyl of galactose seems to be a rate-limiting factor for catalysis by ScGal1p

Another criterion for defining the kinase conformation is the distance between terminal phosphate of ATP and anomeric hydroxyl group of galactose. In *Sc*Gal1p, the distance between terminal phosphate of ATP and galactose varied above and below 6Å (see Figure 4, Panel B). Surprisingly, in *Sc*Gal3p, the distance between ATP and anomeric hydroxyl group of galactose was well below 6Å (see Figure 4, Panel B). Despite this favourable distance for catalysis, *Sc*Gal3p doesn’t catalyse the phosphorylation of galactose because of the distance between D209 and anomeric hydroxyl group of galactose is incompatible with catalysis (see Figure 4, Panel A). On the other hand, in *Sc*Gal3-SAp, the distance between ATP and anomeric hydroxyl group of galactose always remained above 6Å. In fact this distance was highest in *Sc*Gal3-SAp as compared to *Sc*Gal1p and *Sc*Gal3p (see Figure 4, Panel B). This could be a reason why *Sc*Gal3-SAp can’t phosphorylate efficiently [39] despite the distance between D211 and anomeric hydroxyl group of galactose being favourable for catalysis, as the distance between ATP and anomeric hydroxyl group of galactose is not compatible. From these results, it appears that the distance between anomeric hydroxyl group of galactose and D217 is more critical than the distance between terminal phosphate of ATP and anomeric hydroxyl group of galactose. This is based on the observation that addition of SA dipeptide confers kinase activity to *Sc*Gal3p of the order of 1/400 of *Sc*Gal1p [39]. That is, though inefficient, *Sc*Gal3-SAp is nonetheless a kinase, and its kinase activity could be attributed to the H-bond that subsequently forms between anomeric hydroxyl group of galactose and D211.

### H-bond between Y435 and Y57 seems to be responsible for transitioning of *Sc*Gal3-SAp to kinase state

It is interesting to note that although *Sc*Gal3-SAp attains a conformation in which H-bond interaction occurs between D211 and anomeric hydroxyl group of galactose, unlike *Sc*Gal1p, K260 does not form H-bond with D211 in *Sc*Gal3-SAp. In fact, just like *Sc*Gal3p, K260 in *Sc*Gal3-SAp also seems to be moving away from the active site, albeit to a lesser extent. This suggests that there are other structural determinants which play a crucial role independent of K260 in taking *Sc*Gal3-SAp to kinase state. This implies that a second site could be important for regulating the conformation of K266 residue in addition to SA dipeptide. Perhaps, this second site effects the orientation of SA dipeptide as well.

It was previously demonstrated that over-expression of *Sc*Gal3p (i.e. *Sc*Gal3_WT_p) conferred constitutive activation of GAL switch of *S.cerevisiae*. However, over-expression of *Sc*Gal3_Y57W_p did not confer constitutive activation of *GAL* switch [50]. Further, the corresponding substitution Y63W in *Sc*Gal1p abolished kinase activity without interfering in the signalling function [47]. Y63 also happens to be a part of sector 5 (see Table S2). These results highlight the importance of Y63 residue in conferring bifunctionality to *Sc*Gal1p.

Y63 is involved in the binding of galactose in the crystal structure of *Sc*Gal1p. The residue corresponding to Y63 in *Sc*Gal3p is Y57 which is also involved in the binding of galactose in the crystal structure of *Sc*Gal3p. Y63 makes H-bond with Y441 in crystal structure of *Sc*Gal1p. Similarly Y57 (corresponding to Y63 of *Sc*Gal1p, see Figure 3, Panel C) makes H-bond interaction with Y433 (corresponding to Y441 in *Sc*Gal1p, see Figure 3, Panel C) in crystal structure of *Sc*Gal3p. To probe the importance of this H-bond interaction, the distance between Y441 (i.e. the hydroxyl atom of Y441) and Y63 (the hydroxyl atom of Y63) in *Sc*Gal1p, Y435 (the hydroxyl atom of Y435) and Y57 (the hydroxyl atom of Y57) in *Sc*Gal3-SAp and Y433 (the hydroxyl atom of Y435) and Y57 (the hydroxyl atom of Y57) in *Sc*Gal3p was monitored during the above mentioned 150 ns simulation. Y435 and Y57 in *Sc*Gal3-SAp (see Figure 3, Panel C) correspond to Y441 and Y63 of *Sc*Gal1p respectively.

In *Sc*Gal3p it was observed that this H-bond interaction though initially present, breaks as K258 moves away from D209 (see Figure 5 and Figure S4, Panel A & B). In case of *Sc*Gal3-SAp this H-bond (i.e. H-bond between Y57 (corresponding to Y63 of ScGal1p) and Y435(corresponding to Y441 of ScGal1p) forms after D211 starts making H-bond interaction with anomeric hydroxyl group of galactose (see Figure 5 and Figure S2, Panel A & B). Thus, it appears that formation of H-bond between Y435 and Y57 in *Sc*Gal3-SAp is responsible for the transition of *Sc*Gal3-SAp to kinase state. On the contrary, in *Sc*Gal1p, this H-bond is always present (see Figure 5 and Figure S3). Recall that D217 always forms H-bond with anomeric hydroxyl group of galactose in *Sc*Gal1p (see Figure 4, Panel A). Based on this observation, we inferred that H-bond between Y441 and Y63 in *Sc*Gal1p might also be important for its kinase function.

**Figure 5:**
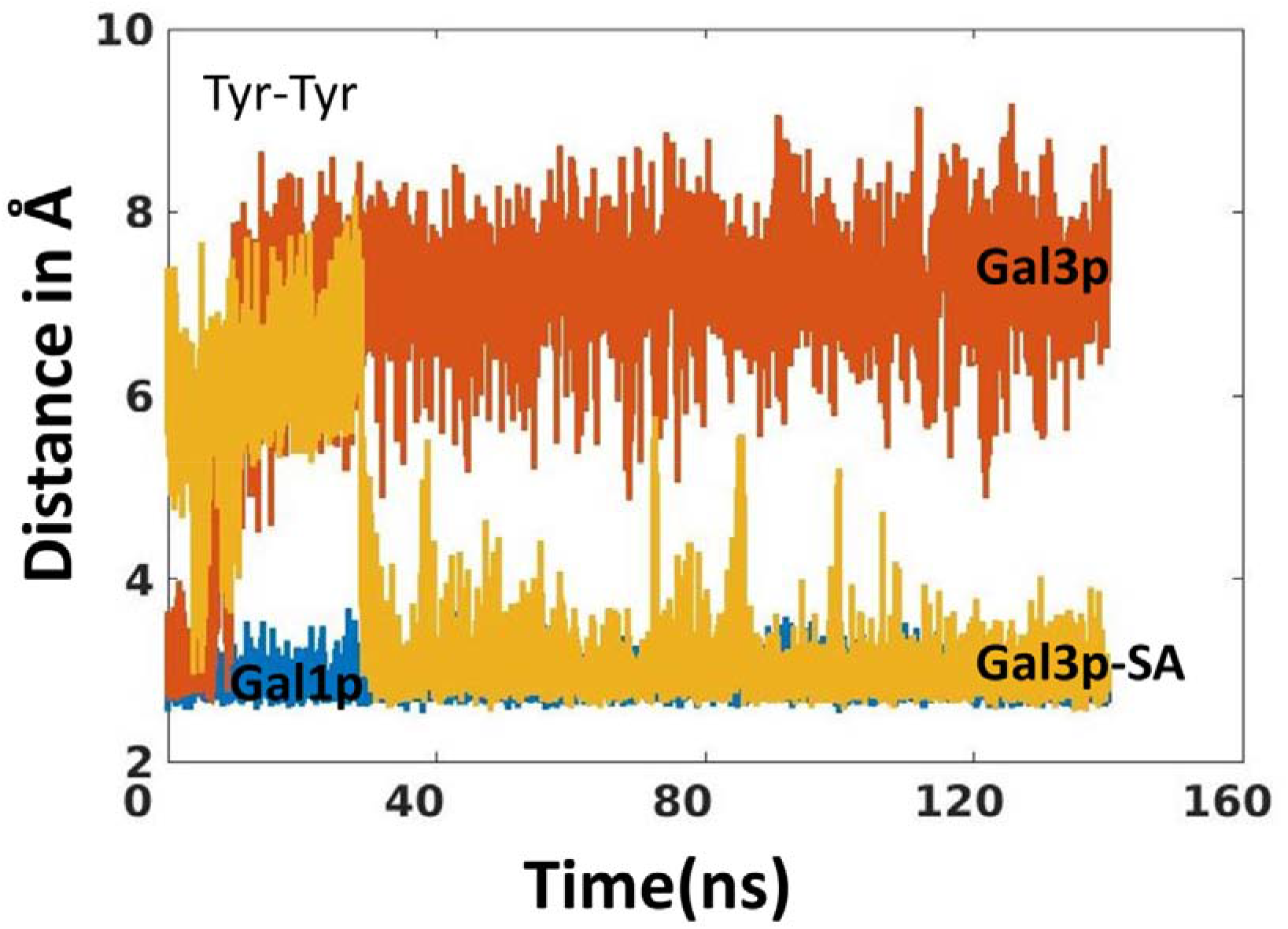
Distance of H bond between Tyr-Tyr in *Sc*Gal1p, *Sc*Gal3p and *Sc*Gal3p-SAp as a function of time. The numbering of galactose binding tyrosine in *Sc*Gal1p isY63 and the interacting tyrosine is Y441. The corresponding Tyr residues distances in *Sc*Gal3p and *Sc*Gal3-SAp are indicated (see figure 3, panel C for numbering of tyrosine residues in *Sc*Gal1p,*Sc*Gal3p and *Sc*Gal3-SAp)

### Conformational sampling was used to bin *Sc*Gal1p into 4 conformational states

From the simulation studies carried out for 150 ns, it was inferred that K266 and Y441 might be important for transition of *Sc*Gal1p to kinase conformation. A prerequisite of *Sc*Gal1p to function as a signal transducer is that it should be in closed conformation. That is the lip distance between V383 and D106 should be less than 35Å (see Figure 2 and corresponding text section). Thus, when *Sc*Gal1p binds to either galactose or ATP in a conformation not suitable for catalysis, provided that it remains in closed conformation, then there is a high probability that *Sc*Gal1p would function as a signal transducer. This is because, catalysis of galactose to galactose-1-phopshate and ADP results in disassociation of galactose-1-phosphate and ADP from *Sc*Gal1p resulting in the protein attaining the open conformation which can’t bind to *Sc*Gal80p. Thus, binding of galactose and/or ATP to *Sc*Gal1p in catalytically incompetent conformation provides *Sc*Gal1p with more leverage to bind to *Sc*Gal80p.

The catalytically incompetent (non-kinase) conformation of *Sc*Gal1p during the period of 150 ns simulation was defined by the distance between terminal phosphate of ATP and anomeric hydroxyl group of galactose being more than 6Å (see Figure 4, Panel B). This is in contrast to *Sc*Gal3p and *Sc*Gal3-SAp, where the lack of H-bond between catalytic aspartate and anomeric hydroxyl group of galactose underlies the lack of kinase activity in *Sc*Gal3p and non-kinase conformation of *Sc*Gal3-SAp. In case of *Sc*Gal1p, distance between anomeric hydroxyl group of galactose and catalytic aspartate always remained within the H-bond range (see Figure 4, Panel A & B). Thus, in order to ascertain, whether the distance between _γ_-phosphate of ATP and anomeric hydroxyl group of galactose is the only parameter that defines the non-kinase conformation of *Sc*Gal1p, we carried out extensive conformational sampling of *Sc*Gal1p by seeding additional 19 trajectories of 150ns each. This was carried out to investigate how often*Sc*Gal1p goes into non-kinase state and if it does, how do the order parameter that defines kinase conformation vary.The order parameter used for discriminating various conformational state was distances between the following functional groups (i) distance between terminal phosphate of ATP and anomeric -OH of galactose. (ii) Distance between anomeric -OH of galactose and D217 (iii) lip distance as measured by the distance between V383 and D108. Based on these parameters, the conformations can be binned into 8 conformational states (see Table. 2, Figure S5, S6 and S7). It is clear that in open conformation there are two subclasses of conformation, one kinase and the other non-kinase. Similarly, in the closed conformation, we have kinase and non-kinase (Figure 6).

**Figure 6:**
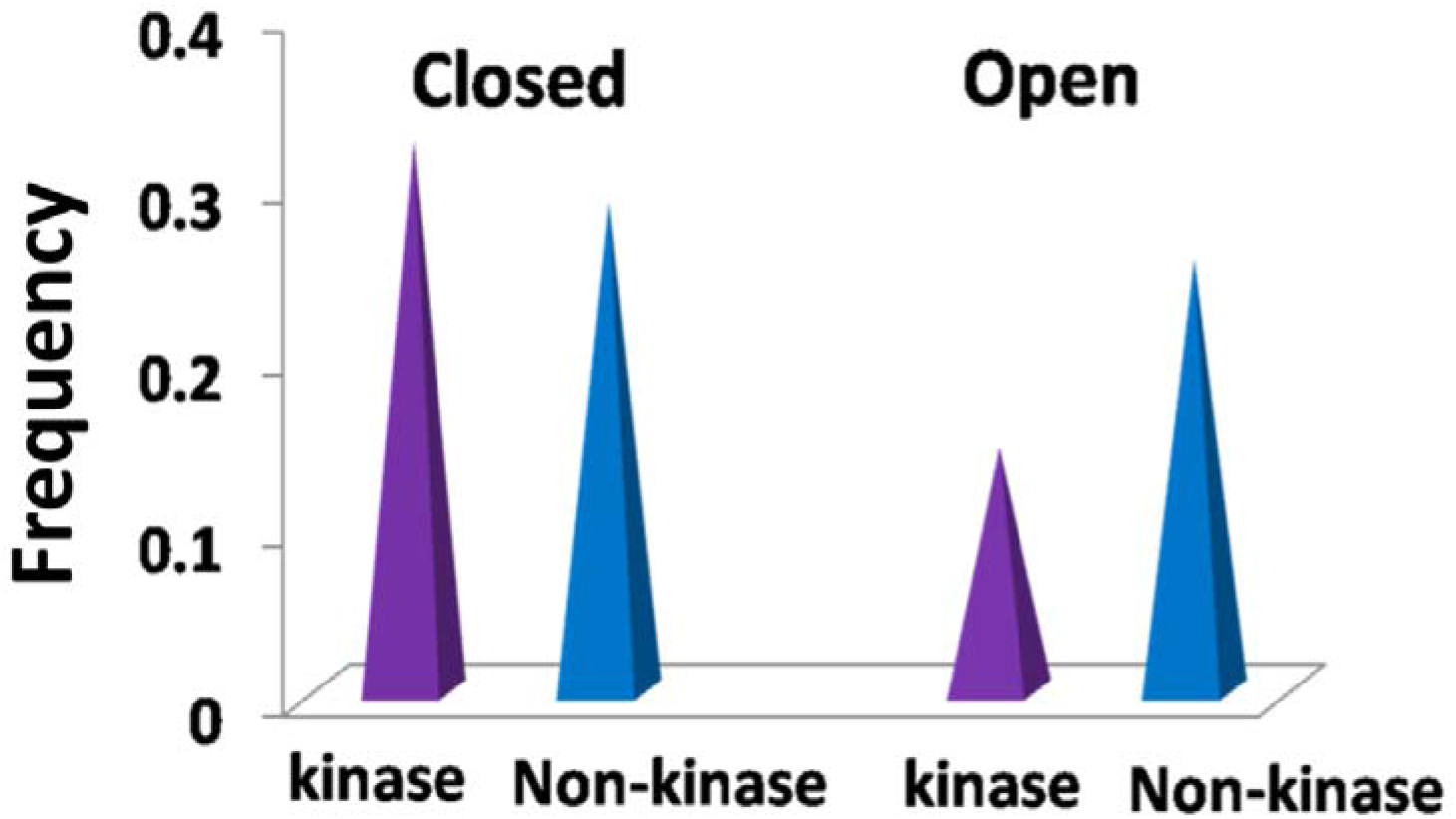
Relative proportion of kinase and non-kinase conformations in open and closed states of *Sc*Gal1p. Given that Y63, S171, D217, K266 andY441 are important for transition of *Sc*Gal1p to kinase state, we decided to look into the distribution of (i) distance between K266 and D217. (ii) Distance between K266 and S171 in kinase and non-kinase conformation (Figure 7A & B). From this analysis we infer that as K266 moves closer to D217 that is away from S171 the protein acquires kinase conformation (Figure7 and figure S8). That is, as long as K266 is towards S171, it remains in non-kinase state (Figure7 and figure S9A & B).The distribution of distance between Y441 and Y63 remains within the range compatible for H bond interaction through out the simulation. That is, this parameter remains same both in kinase and non-kinase states of *Sc*Gal1p (data not shown).

Thus, in *Sc*Gal1p the transition to signal transducer state is governed by the motion of K266 from D217 to S171. This motion of K266 away from D217 and towards S171 results in the distortion of catalytic loop. In contrast to this, in *Sc*Gal3p, it was observed that K258 not only moves away from D209 and S164 (i.e. the galactose/ATP binding site,) there is also an increase in the distance between Y433 and Y57 to as much as 7Å, resulting in a decoupling of galactose from the catalytic loop (Figure S9A, B & C). This happens to be the major difference in the signal transducer conformation of *Sc*Gal1p and *Sc*Gal3p.Given that our studies implicate a role for S171 in governing the motion of K266 in *Sc*Gal1p, a possibility could be that in case of *Sc*Gal3p, absence of SA dipeptide results in K258 undergoing unhindered motion. This unhindered motion of K258 could potentially result in an increase of distance between Y433 and Y57 to as much as 7Å, resulting in decoupling of galactose from the catalytic loop. Based on the above considerations, we conjecture that the above sequence of events permanently freezes*Sc*Gal3p in signal transducer state from where it cannot transition to kinase state.A corollary of this observation is that, if the H bond between Y441 and Y63 is broken, *Sc*Gal1p would have more signalling activity and less kinase activity.

If the interaction between Y441 and Y63 is functionally relevant, why Y441 is neither a conserved residue nor is coevolving with Y63 which is a part of sector 5.For example,*Zygosaccharomyces bailii,* a species that diverged before whole genome duplication [53] has two sequences similar to *GAL1/GAL3* of *S.cerevisaie* (NCBI). In both these sequences, the residue corresponding to Y441 is cysteine which can’t form H bond with the residue corresponding to Y63. If the corresponding residue to Y441 is substituted by cysteine in *Zygosaccharomyces bailii* by chance and has no biological implication, then substitution of Y441 in *Sc*Gal1p is not expected to alter the kinase and/or signaling function. Based on this, we wanted to know, what would happen if Y441 is replaced by alanine in *Sc*Gal1p. The second residue we picked for mutational analysis was K266. Galactokinases have either Lys or Arg at the catalytic site [54]. It is intriguing that why some galactokinases use Lys while other galactokinase use Arg (see discussion for more details). We were therefore interested in knowing the consequences of substituting K266 with arginine.

### Phenotypic analysis indicated that *Sc*GAL1_K266R_pand *Sc*GAL1_Y441A_p variants gained signalling activity

We introduced K266R and Y441A substitutions using SDM and integrated the *GAL1_K266R_* and *GAL1_Y441A_* alleles at the *GAL1* locus of *Sc723 gal3Δ GAL1_wt_* (see material and methods for details) to generate *Sc723gal3ΔGAL1_Y441A_* and *Sc723 gal3ΔGAL1_K266R_* strains. It is reported a *gal3Δ* strain shows long term adaptation phenotype [27, 34, 55]. Long term adaptation phenotype can be demonstrated by spotting assay [56, 57]. Using a plating assay for colony forming units, [57] had demonstrated that a population of *Sc723gal3ΔGAL1_wt_* when exposed to galactose, only 3 cells out of 1000 respond to galactose and such cells, referred to as rapid inducers, eventually populate the culture ensuing LTA. Consistent with the above, increasing the basal expression of *GAL1* by inactivating the antisense *GAL10* ncRNA, increased the rapid inducer count by 7 fold [58]. If *Sc*Gal1_K266R_p and *Sc*GAL1_Y441A_p variants had acquired altered signalling properties, we expected to observe a change in the phenotype as well as the number of rapid inducers with reference to *Sc723gal3ΔGAL_wt_* strain.

We first carried out a phenotypic analysis of the above strains to determine whether the *gal3Δ* strains carrying mutant *GAL1* alleles show any discernible difference from the *gal3Δ* carrying the wild type *GAL1*. Strains of indicated genetic background grow equally well on complete medium with glycerol plus lactate as the sole carbon source (Figure 8A). Growth pattern of the same strains on medium supplemented with 2.0% galactose containing 0.05% ethidium bromide is indicated in Figure 8, Panel B. Supplementation of ethidium bromide was required to abolish the background growth as reported previously [56,57]. Under the above experimental conditions, as expected *Sc723gal3Δgal1Δ* strain (negative control) does not grow while *Sc723GAL3_wt_ GAL1_wt_* strain (positive control) grows. On the other hand, *Sc723gal3ΔGAL1_wt_* grows better than the negative control as expected. As compared to the positive control, the growth phenotype of *Sc723 gal3Δ GAL1_wt_* is clearly retarded and exhibits LTA. It is clear that Sc723*gal3ΔGAL1_K266R_* does not grow on this medium. This indicates that *Sc*Gal1_K266R_p variant is either completely devoid of kinase activity or signalling or both. It is also possible that both these activities are reduced to a level that is insufficient to show growth on this medium. Alternately, it is possible that this substitution resulted in the loss of stability of the protein product, resulting in the above phenotype. On the other hand, *Sc*Gal1_Y441A_p variant exhibits marginally better growth than *Sc723 gal3Δ GAL1_wt_*.

**Figure 7:**
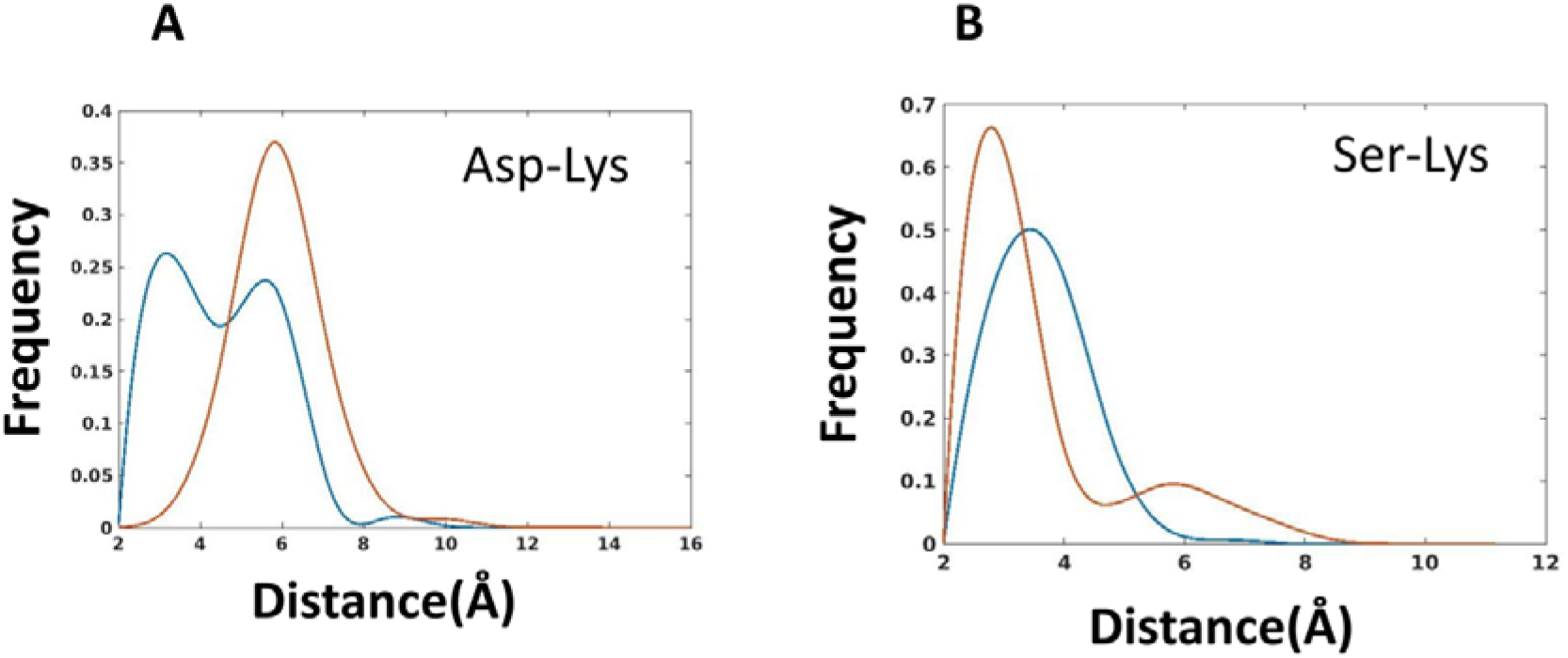
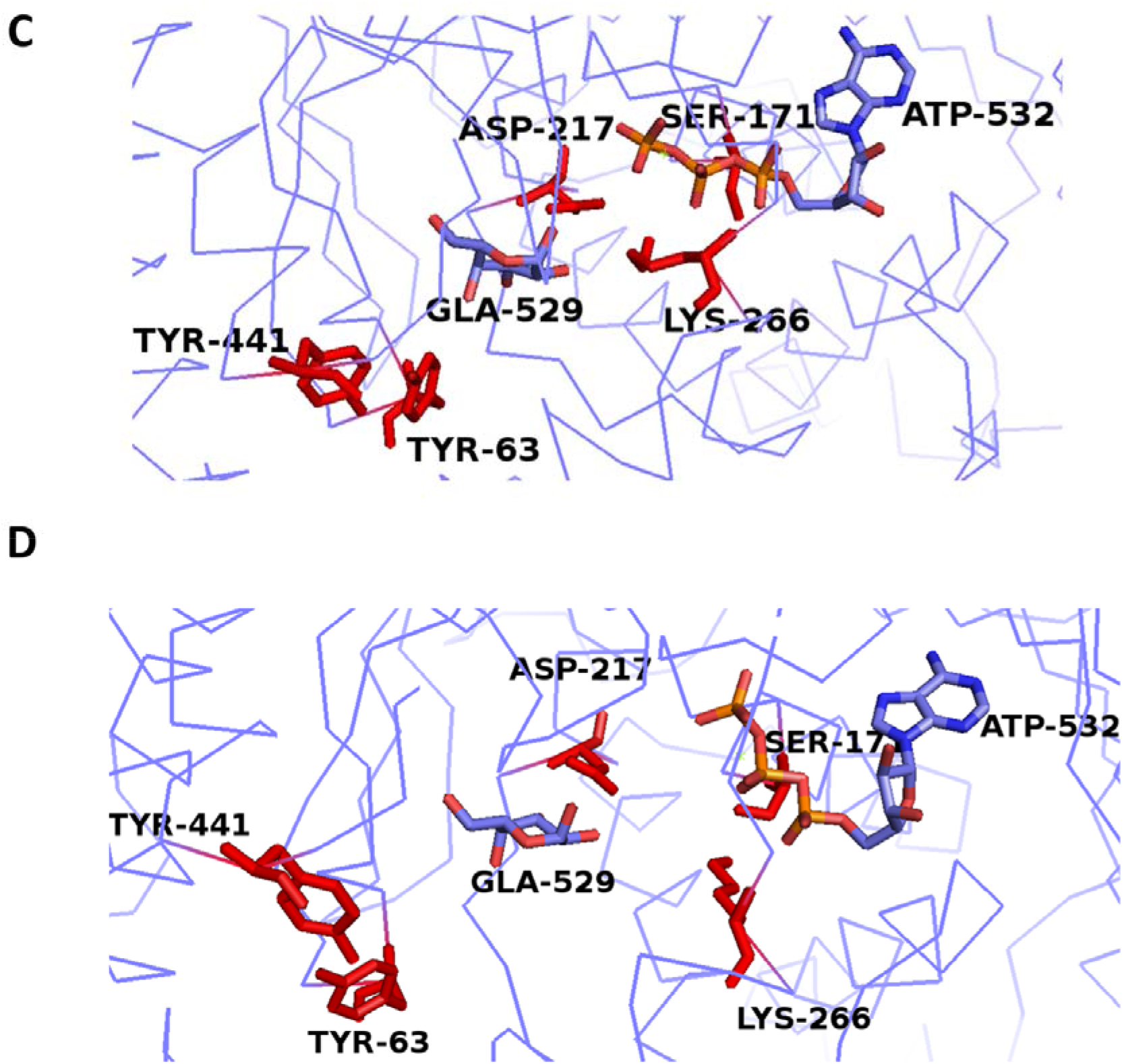
Orientation of K266 and Y441 in kinase and non-kinase conformations of *Sc*Gal1p. Panel A shows the distribution of distances between K266 and D217 in kinase and non-kinase states of *Sc*Gal1p. The blue curve corresponds to kinase state and red curve corresponds to non-kinase state. Panel B shows the distribution of distances between K266 and S171 in kinase and non-kinase states of *Sc*Gal1p. The blue curve corresponds to kinase state and red curve corresponds to non-kinase state. Panel C and D show the structural orientation of Y63,S171,D217,K266,Y441,galactose and ATP in kinase and non-kinase states respectively.

**Figure 8:**
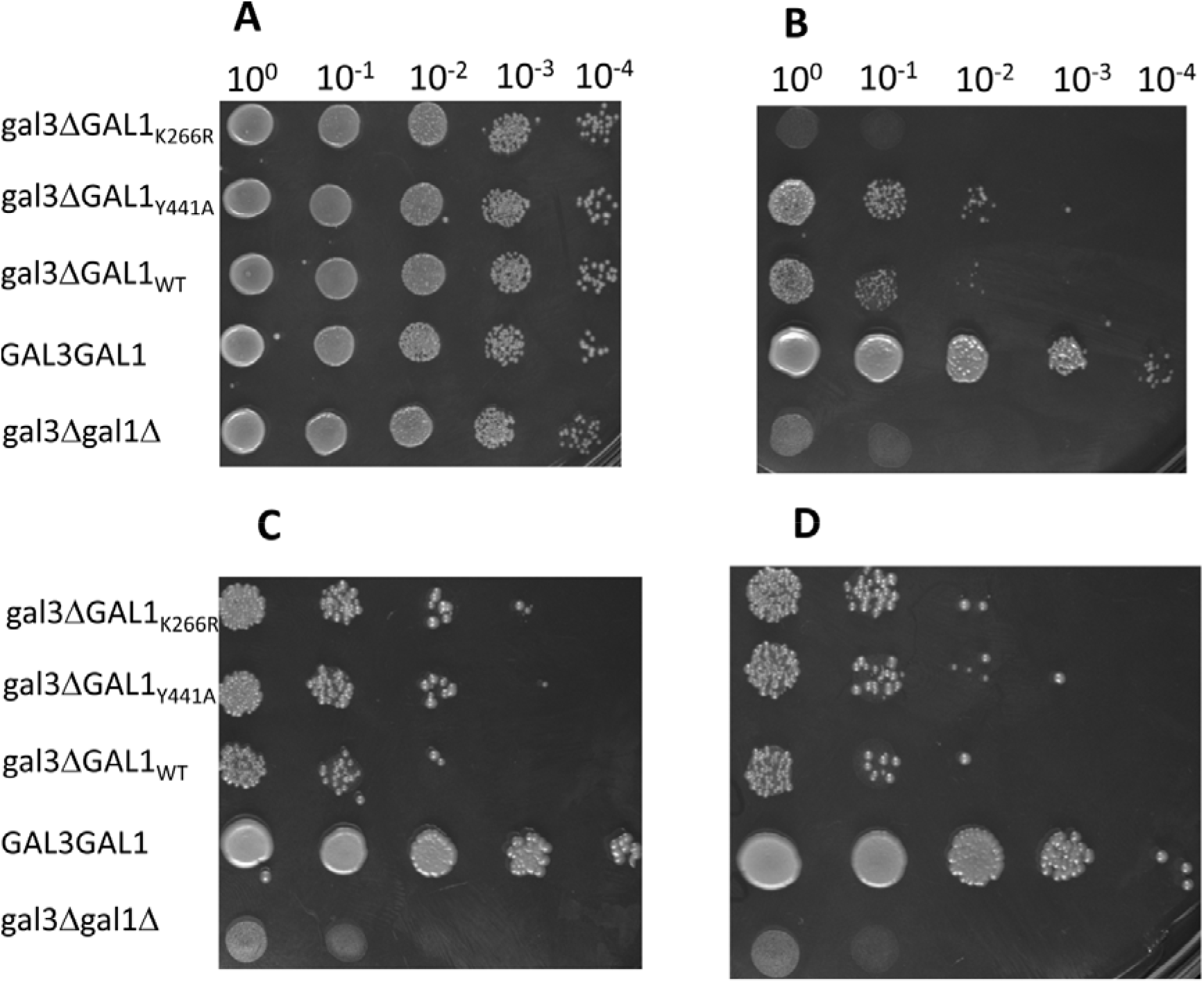
Phenotypic analysis of strains bearing wild-type and mutant derivatives of *GAL1* in a gal3Δ background. Panel A show the growth of 20µl of cell suspension appropriately diluted from O/N culture of indicated strains pre-grown in complete glycerol plus lactate medium and spotted on complete media with glycerol plus lactate as the sole carbon source. gal3ΔGAL1_K266R_ refers to *Sc723gal3*Δ*GAL1_K266R_*. gal3ΔGAL1_Y441A_ refers to *Sc723gal3*Δ*GAL1_Y441A_*. gal3ΔGAL1_WT_ refers to *Sc723gal3*Δ*GAL1_WT_*. gal3Δgal1Δ refers to *Sc723gal3*Δ*gal1*Δ. GAL3GAL1 refers to *Sc723GAL1_WT_GAL3_WT_ Sc723gal3*Δ*gal1*Δ and *ScGAL3_wt_GAL1_wt_* serve as the negative and positive control respectively. *Sc723gal3*Δ*GAL1_wt_* strain is taken as the reference for comparing the behaviour of the GAL1 variants. Panel B is same as in Panel A, exc pt the sole carbon source is 2% galactose along with ethidium bromide. Panel C and D is same as panel A, except that the medium consists of glycerol plus lactate histidine drop out with 0.05% and 0.5% galactose respectively.

Similar phenotypic analysis was also carried out using *ScHIS3* as a reporter. Here, a *GAL1p::HIS3* cassette is integrated at the *URA3* locus, in the above strains, otherwise deleted for *HIS3* gene [41, 57]. Therefore, these strains can’t grow on histidine drop out medium with glycerol plus lactate as the carbon source, unless the *ScHIS3* expression is driven by the activation of *GAL1* promoter. In this assay, galactose is added as an inducer. Therefore, a strain with a kinase minus function should show growth, provided it has retained the signalling function. Both positive and negative controls show the expected phenotype. The results clearly show that both *Sc*GAL1_K266R_p and *Sc*GAL1_Y441A_p variants have increased signalling activity as compared to *Sc*Gal1_wt_p at 0.05% as well as 0.5% concentrations of galactose(Figure 8 Panel C and D). Based on this result, it is inferred that*Sc*Gal1_K266R_p variant may be devoid of galactokinase activity but not signalling.

To determine whether *Sc*Gal1_K266R_p and *Sc*Gal1_Y441A_p variants have acquired constitutive activity, reporter assay was performed in which galactose was not added as the inducer. (Figure9, Panel A and B). It is clear from Figure9, that, only *Sc723gal3ΔGAL1_K266R_* gave rise to distinct colonies. Note that, out of 10^5^ cells spotted, only a tiny fraction of cells gave rise to distinct colonies. Neither *Sc723gal3ΔGAL1_wt_* nor *Sc723gal3ΔGAL1_Y441A_* strains gave rise to distinct colonies.

We considered that these colonies could arise either because of recessive mutations in *ScGAL80* or because of dominant mutations in *ScGAL4*. Either of these mutations would render the strains constitutive [59], in that, the *GAL* switch is induced even in the absence of galactose. As we did not observe such colonies in *Sc723gal3ΔGAL1_wt_* and*Sc723gal3ΔGAL1_Y441A_*, the colonies observed in *Sc723gal3ΔGAL1_K266R_* are unlikely to bear dominant constitutive mutations. Nevertheless, to ascertain whether these colonies carry mutations described above or whether it is a physiological response, following experiment was carried out.

Cells from two independent colonies of *Sc723gal3ΔGAL1_K266R_* were resuspended separately in complete liquid medium with glucose or with glycerol plus lactate as the carbon source. These cultures were allowed to grow for at least ten generations. If these colonies were to bear constitutive mutations as indicated above, all the descendants should bear this mutation regardless of the carbon source used for their growth. Spotting these cells pre-grown in glucose or glycerol plus lactate should grow on histidine drop out medium, provided these cells bear any of the above mutations that cause constitutive activation. On the other hand, if they are physiological revertants, then growing in glucose for ten generations would wipe out that physiological state due to glucose repression. However, growing them in glycerol plus lactate would not wipe out this state, as it is not a repressing carbon source. That is, in glycerol plus lactate medium, once the *GAL* switch is on, it would sustain itself and remain induced for *GAL* genes indefinitely (as long as glucose is not present) due to positive feedback loop of *GAL1*. As can be seen from the results,(Figure 9, panel C, D and E) cells pre-grown on glucose for ten generation upon spotting again showed the original phenotype but not when cells are pre-grown on glycerol lactate. Based on this, we infer that *Sc*Gal1_K266R_p variant has constitutive activity.

**Figure 9:**
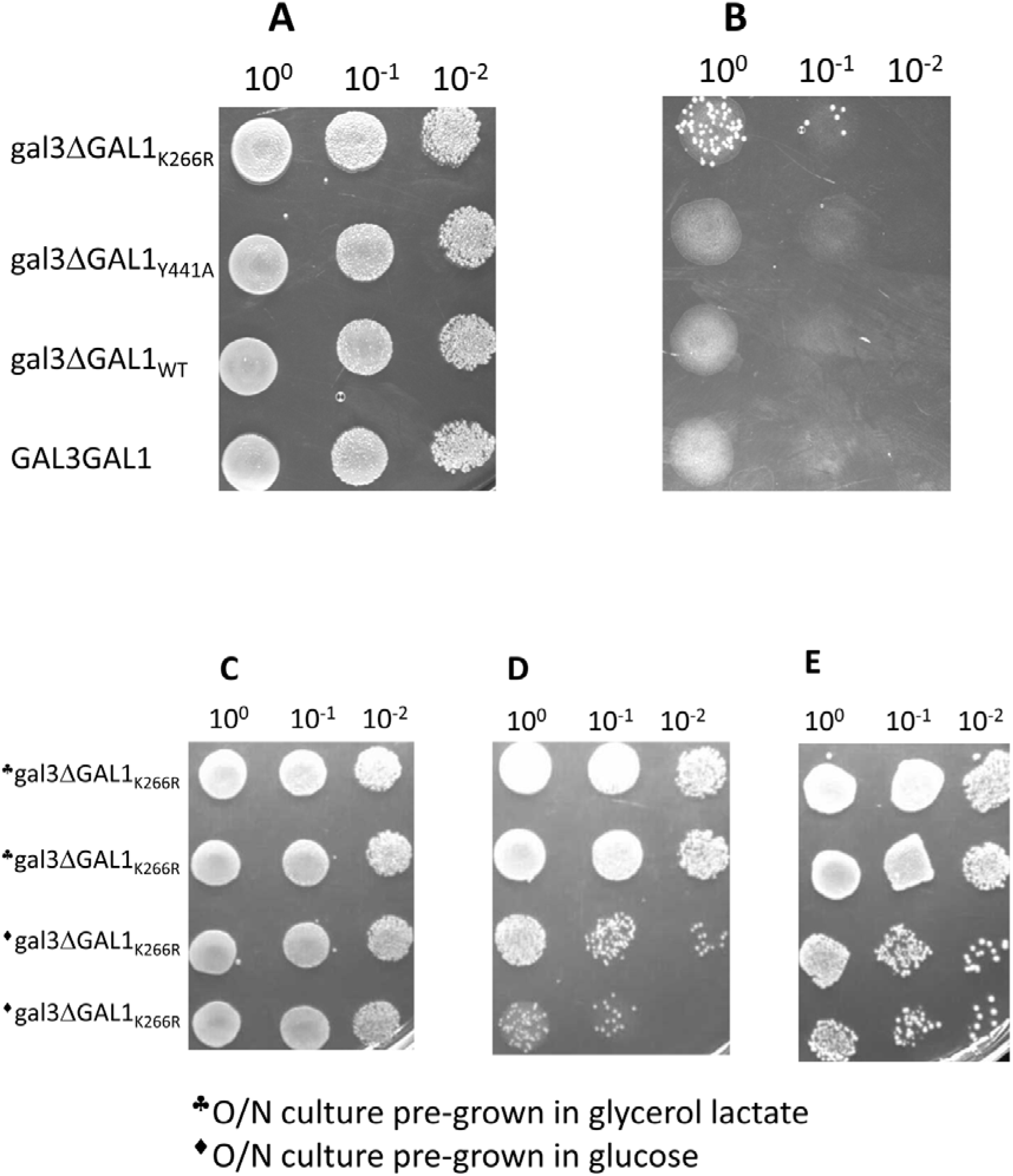
Phenotypic analysis of presence of constitutivity in mutant derivatives of *Sc*Gal1p in gal3Δ background. Panel A shows the growth of 50µl of cell suspension appropriately diluted from O/N culture of indicated strains pre-grown in glycerol lactate medium and spotted on complete media with glycerol lactate medium. Panel B is same as panel A except that the medium is Histidine drop out <u>without</u> galactose. Cells from two independent colonies (see panel B) were collected and grown in complete glycerol and glucose medium separately for at least 10 generations. 50µls of cells from these pre-grown cultures were spotted on complete glycerol lactate medium (panel C) histidine drop out glycerol plus lactate plates without galactose (panel D) or histidine drop out glycerol plus lactate plates with 0.5% galactose (Panel E). ♣ refers to O/N culture pre-grown in glycerol plus lactate. ♦ refers to O/N culture pre-grown in glucose.

If *Sc*GAL1_K266R_p variant can activate the switch independent of galactose (that is, it is constitutive), why did only few out of 10^5^ cells give rise to distinct colonies (Figure 9 Panel B). This non-genetic heterogeneity in the constitutive phenotype is most likely driven by the stochastic expression *Sc*GAL1_K266R_p, (note that the *ScGAL1_K266R_* allele is in a *gal3Δ* background), thus rendering only few colonies of *Sc723 gal3ΔGAL1_K266R_* to appear under the above experimental conditions (See discussion for more details).

### Sc723gal3ΔGAL1_K266R_ and Sc723 gal3ΔGAL1_Y441A_ give rise to more rapid inducers than Sc723gal3ΔGAL1_wt_

The number of rapid inducers were quantified as described in material methods using two growth conditions as reported previously [57]. When rapid inducers were measured using galactose as a carbon source, *Sc723gal3ΔGAL1_Y441A_* showed close to 6 fold increase in rapid inducers (Table 3). This technique could not be used for determining rapid inducers for *Sc723gal3ΔGAL1_K266R_*, as it does not grow under the conditions of the experiment (see figure 8 B). When rapid inducers were quantified using growth on histidine drop out medium, both *Sc723gal3ΔGAL1_Y441A_* and *Sc723gal3ΔGAL1_K266R_* showed a four to five fold increase in rapid inducers as compared to *Sc723gal3ΔGAL1_wt_* (Table 3).

**Table 2:**
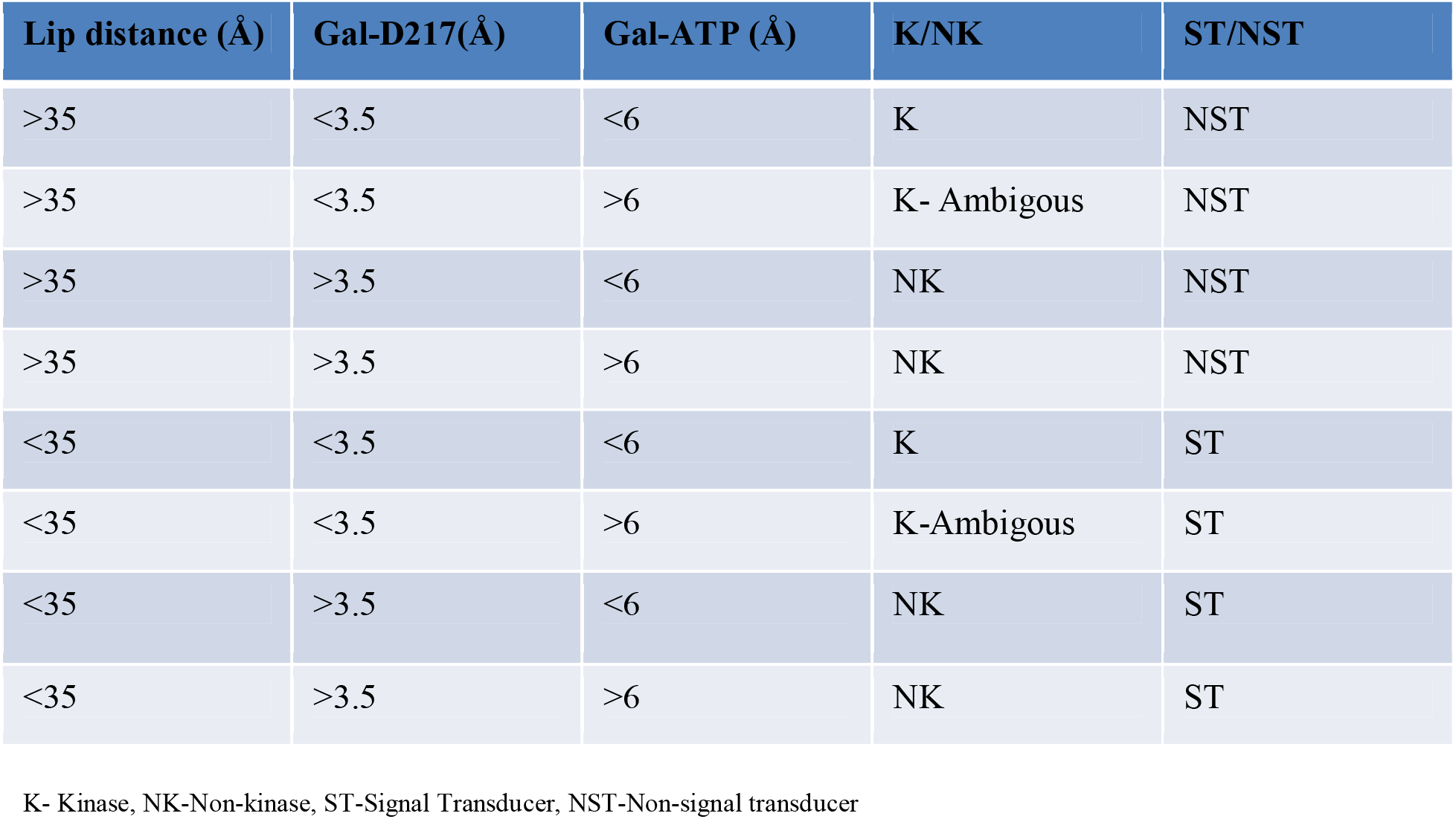
Classification of 8 functional states of ScGal1p as Kinase and Signal transducer.

**Table 3:**
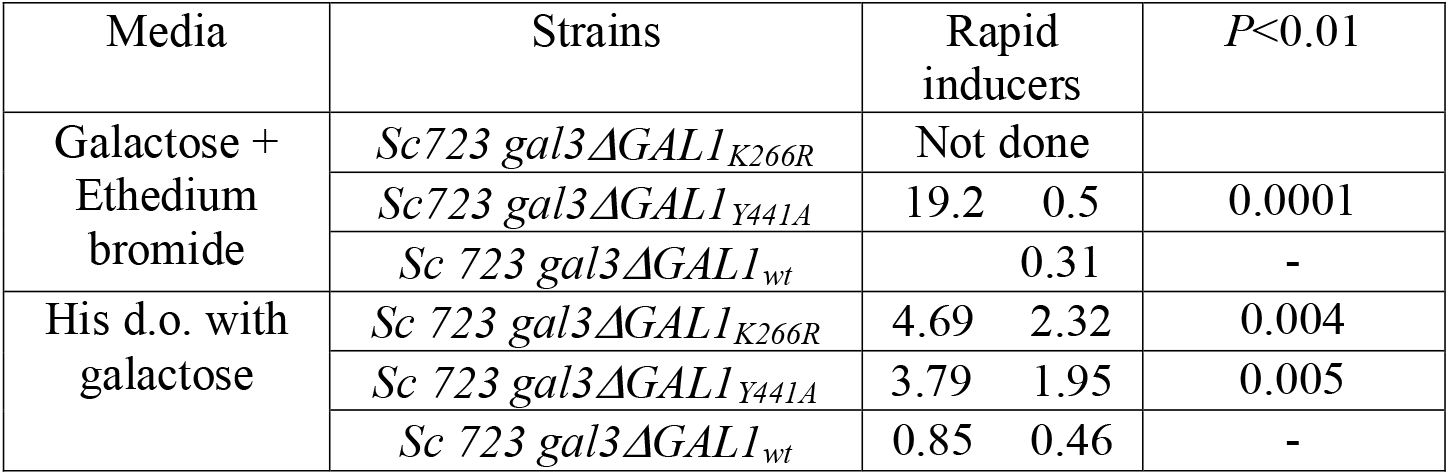
Rapid inducer assay.

### Fraction of ON cells in Sc723gal3ΔGAL1_K266R_ and Sc723gal3ΔGAL1_Y441A_ appear earlier than *Sc723gal3ΔGAL1_wt_*

We carried out FACS analysis to determine the fraction of ON cells in response to 0.5% galactose.*Sc723GAL3_wt_GAL1_wt_*with integrated *GAL1p::GFP*cassette at the *URA3* locus was available in the laboratory [57] and served as the positive control. We integrated *GAL1p::GFP* cassette at the *URA3* locus in *Sc723gal3ΔGAL1_wt_*, *Sc723gal3ΔGAL1_Y441A_* and *Sc723gal3ΔGAL1_K266R_* as described in material and methods. The autofluoresence of *Sc723 gal3Δ GAL1_wt_* with an integrated copy of *GAL1p:GFP* at the *URA3* locus grown in complete glycerol plus lactate media was used to draw the cut off for determining the fraction of ON cells. Based on this, cells with fluorescence intensity above 10^2^ were considered ON cells. Above strains were pre-grown in complete glycerol plus lactate liquid medium upto mid-log phase. These cells were inoculated into histidine drop out glycerol plus lactate medium containing 0.5% galactose, such that the initial cell concentration was 10^6^cells/ml. Fifty thousand cells collected at the indicated time points were subjected to FACS analysis to determine the fraction of ON cells (see materials and methods for details).

It is clear from this analysis that by 18 hours post inoculation, the fraction of GFP expressing cells of *Sc723GAL3_wt_GAL1_wt_* had reached 100%. As expected the fraction of GFP expressing cells in *Sc723gal3ΔGAL1_wt_*, started increasing by 52 hrs and it took nearly 66 hours to reach a state where nearly 100% of the cells were GFP expressing. In contrast, the fraction of GFP expressing cells in *Sc723gal3ΔGAL1_Y441A_* started increasing by 44 hours. By 56 hours the cells were fully induced. The range of intensity of GFP expression remained the same between all the three strains (See figure10A, Figure S10A and B).

In contrast to the above, the behaviour of *Sc723gal3ΔGAL1_K266R_* was different from *Sc723 gal3ΔGAL1_Y441A_* and *Sc723gal3ΔGAL1_wt_*. First, even at the 0 time point, *Sc723gal3Δ GAL1_K266R_* had higher fraction of induced cells (Figure 10A and FigureS10A). This is consistent with the phenotype reported in Figure 9. This number started increasing much before *Sc723gal3ΔGAL1_wt_*. The second difference was that the maximum intensity of induction was clearly different than what is observed in *Sc723GAL3_wt_GAL1_wt_*, *Sc723gal3ΔGAL1_wt_* or *Sc723gal3ΔGAL1_Y441A_*(Figure S10A). This increase was of the order of five fold. Third, *Sc723gal3ΔGAL1_K266R_* gave higher fraction of ON cells till 44 hrs after which its kinetics was similar to *Sc723gal3ΔGAL1_wt_* and not similar to *Sc723gal3ΔGAL1_Y441A_*. These differences in the induction kinetics probably reflect a difference in the binding ability of these variants with galactose.

**Figure 10:**
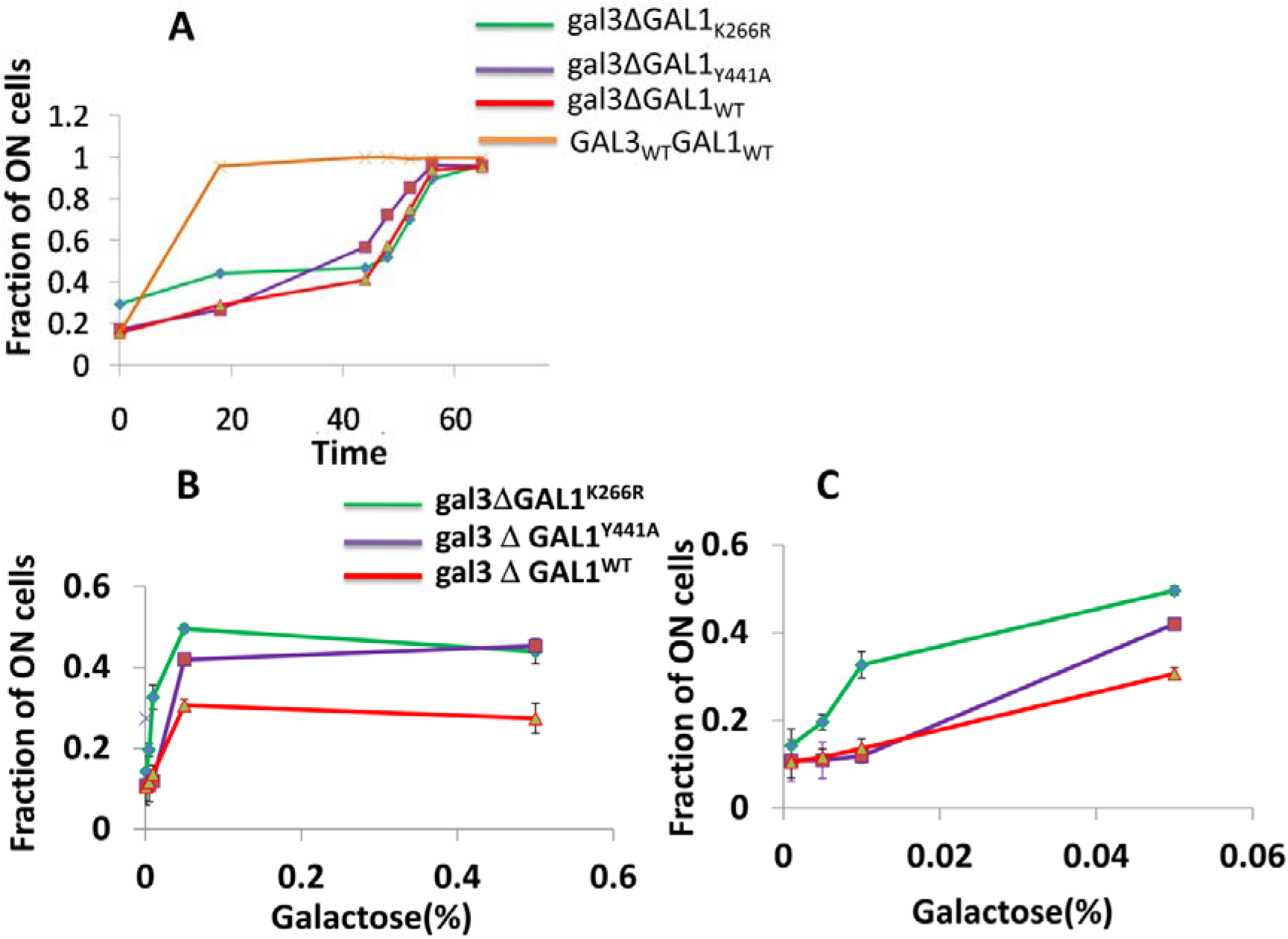
Fraction of ON cells in *Sc723Δgal3GAL1_WT_,*Sc723gal3*ΔGAL1_K266R_* and *Sc723gal3ΔGAL1_Y441A_* in response to galactose. Panel A shows the kinetics of *GAL1* promoter driven GFP expression in response to 0.5% galactose in indicated strains. Panel B shows the response of indicated strains to increasing concentration of galactose(0.001%-0.5%). Panel C is a replot of panel B with the concentration of galactose is shown up to only 0.05% of galactose. This is to highlight the difference in the induction between the indicated strains for lower concentration of galactose.

To test the above possibility we carried out a dose response analysis by measuring fraction of ON cells in*Sc723gal3ΔGAL1_wt_*, *Sc723gal3ΔGAL1_K266R_* and *Sc723 gal3ΔGAL1_Y441A_*at the end of 42 hours of induction in response to galactose concentration ranging from 0.001%to 0.5% of galactose (Figure 10B & C). It is observed that both *Sc723gal3ΔGAL1_K266R_* and *Sc723gal3ΔGAL1_Y441A_* showed increased GFP expression at concentration of galactose much less than what is required for the*Sc723gal3ΔGAL1_WT_*. However, the concentration of galactose required to induce GFP in *Sc723gal3ΔGAL1_K266R_* was lower than what was required for *Sc723gal3ΔGAL1_Y441A_* (Figure 10B & C and Figure S10C & D). Thus, it appears that *Sc*Gal1_K266R_p variant has a higher sensitivity towards galactose than *Sc*Gal1_wt_p and *Sc*Gal1_Y441A_p. However, this property alone may not be sufficient to explain the differences in growth kinetics observed above. This is because, GFP expression induced by galactose involves multiple equilibria and also depends on the expression levels of the wild type and the mutant versions of *Sc*Gal1p.

### Galactokinase activity and steady state expression of*Sc*Gal1_K266R_p and *Sc*Gal1_Y441A_p variantsis different from *Sc*Gal1_wt_p

To determine whether the differential phenotypic response of *Sc*Gal1_K266R_p and *Sc*Gal1_Y441A_p variants is due to the mutations *per se* or because of a secondary consequence of a difference in the steady state expression, we monitored the steady state expression of the *Sc*Gal1_WT_p and the two protein variants using western blot analysis (see methods for details). To carry out this experiment, *ScGAL80* was deleted in *Sc723gal3ΔGAL1_wt_*, *Sc723gal3ΔGAL1_K266R_* and *Sc723gal3ΔGAL1_Y441A_*(see methods for details), to circumvent the inherent difference that could otherwise be associated with the expression of Gal1p and the variants, because of the possible difference in the autoregulation implemented by these variants. Protein extracts were prepared after these strains were grown to mid-log in complete glycerol plus lactate medium. Protein expression was probed with antisera raised against *Sc*Gal3p [47], assuming that the antibodies recognise the wildtype and the mutant versions to the same extent. The experiment was repeated several times at different protein concentrations and the band intensity was quantified and normalised to G6PD (see Figure11 and Figure S11).

The difference in expression between *Sc*Gal1_wt_p and *Sc*Gal_K266R_p was found to be significant with *P*<0.002 and between *Sc*Gal1_Y441A_p and *Sc*Gal1_WT_p with *P*< 0.02. It is calculated from the above results, that the steady state expression of *Sc*Gal1_K266R_p and *Sc*Gal1_Y441A_p is reduced by 46.9% and 18.7% than wild type respectively (see Figure11, Panel A and figure S11,Panel A).

Specific activity of galactokinase determined in the same cell extracts increased by 13.9% for *Sc723gal3ΔGAL1_Y441A_* as compared to *Sc723gal3ΔGAL1_wt_*(*P<0.0022).* On the other hand, specific activity of galactokinase in *Sc723 gal3ΔGAL1_K266R_*is reduced by 50.8% as compared to *Sc723 gal3ΔGAL1_wt_* with *P*<0.00003 (See figure 11, Panel B).

**Figure 11:**
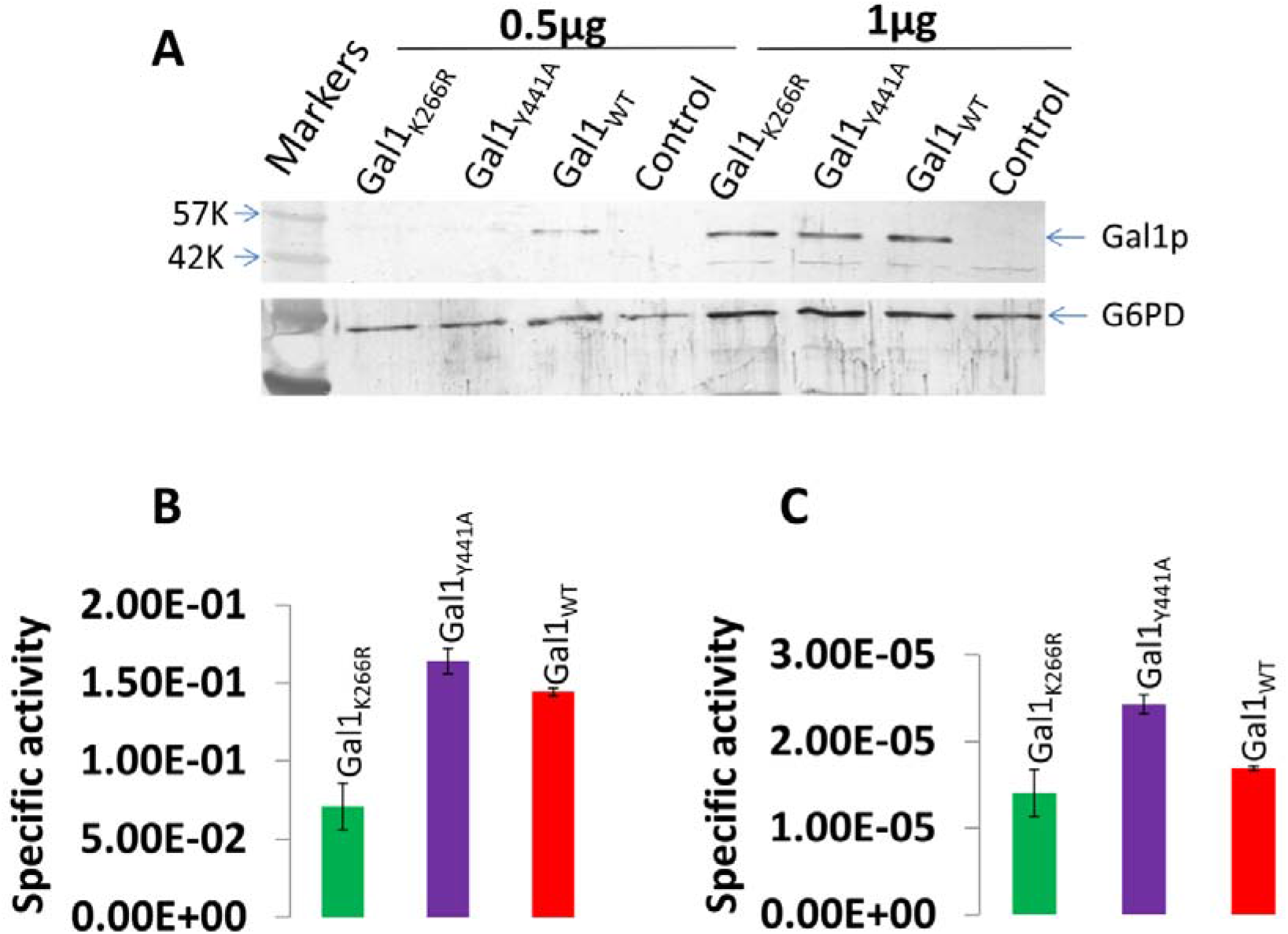
Western blot analysis (panel A) and determination of galactokinase activity of wild type and mutant versions of *Sc*Gal1p. Panel A shows the expression of mutant and wild type Gal1p in *gal80*_Δ_ background. Cell free extract was probed by Gal3 antisera for detection of Gal1p. G-6-P-D antisera was used as a loading control. _Panel B represents nanomoles of galactose 1 phosphate formed per minute per microgram of total protein. Panel C represents the specific activity normalized against the AU of band intensity._

To obtain the galactokinase activity per molecule of the mutant proteins as compared to the wild type, the activity was normalised to the band intensity. This analysis indicated that the activity per molecule of *Sc*Gal1_K266R_p reduced by 16.9% as compared to that of *Sc*Gal1_wt_p(*P*<0.031).On the other hand the activity per molecule of *Sc*Gal1_Y441A_p is increased by 43.8% as compared to that of *Sc*Gal1_wt_p.(*P<*0.00002)(See Figure 11 panel C).

## Discussion

*GAL* genetic switch of *S.cerevisiae* is a model to understand regulation of eukaryotic transcriptional activation, epigenetic transcriptional memory as well as functional diversification following gene duplication. *ScGAL1* and *ScGAL3* evolved by duplication [60] of a bifunctional common ancestor consisting of galactokinase and signalling activity [61]. It is suggested that sub-functionalisation that occurred following duplication of *ScGAL3* and *ScGAL1* was to escape from adaptive conflict that existed in the common ancestor [61]. During sub-functionalisation, *ScGAL3* retained basal expression but fold induction was reduced [61]. At the protein level, *Sc*Gal3p underwent many changes and the deletion of the SA dipeptide is thought to be the last event that abolished the kinase activity completely [39, 62, 63]. Thus, *Sc*Gal3p got completely sub-functionalised at the expression and at the activity level. In contrast, *Sc*Gal1p lost basal expression but gained high fold induction thus getting sub-functionalised at the level of expression. On the other hand, sub-functionalisation of *Sc*Gal1p at the activity level, is inferred to be partial as its signalling function is considerably less as compared to *Sc*Gal3p [26, 46, 61, 63, 64, 65].Sub-functionalisation at the level of expression is based on promoter swap experiments between Sc*GAL3* and *ScGAL1* followed by measuring fitness [61, 63, 64].

Based on the results presented here, we suggest that it is unlikely that *Sc*Gal1p got partially sub-functionalized by losing the signalling activity partially. Comparing *Sc*Gal1p’s signalling activity to *Sc*Gal3p to infer that *Sc*Gal1p is partially sub-functionalised is misleading for the following reason. From the protein conformation-function perspective, a complete loss of kinase activity allows all the functional *Sc*Gal3p molecules to exist only in a conformational state compatible with signalling. As galactokinase exists in two conformational states compatible with the two distinct functions (see below for details), galactokinase is not expected to have as much signalling activity as *Sc*Gal3p, to start with. Further, it can also be interpreted that *Sc*Gal3p acquired more signaling activity than its ancestor, rather than considering that *Sc*Gal1p partially lost the signalling function. Thus, we suggest that to infer whether *Sc*Gal1p or *Sc*Gal3p is sub-functionalised at all or not, signalling activity of these proteins be compared with the signalling activity of its ancestral bifunctional galactokinase and not with each other.

Second, if *Sc*Gal1p is sub-functionalised with respect to signaling activity, it is not clear as to why *Sc*Gal1p only got partially sub-functionalized and not completely sub-functionalized similar to what happened with *Sc*Gal3p. Many substitution mutations that completely knock off signalling function and yet retain the kinase function have been reported (See Table 1). A recent genetic screen for mutations that specifically knock of the transcriptional memory, identified a single substitution of D117V, that abolished signaling function. Strain bearing D117V substitution is normal for *GAL* induction [48]. This suggests that *Sc*Gal1p had the potential to get completely sub-functionalized and yet species that carry such galactokinases have not been identified.

Third, it was reported that the retention of epigenetics transcriptional memory of a previous induction state is due to the signal transduction function of *Sc*Gal1p and not *Sc*Gal3p [48, 66]. These results suggest that the retention of signalling activity by *Sc*Gal1p is unlikely to be due to chance alone. Based on all of the above, we suggest that the most plausible events that would have occurred after duplication of the ancestral bifunctional galactokinase, is the *Sc*Gal1p did not loose any signalling activity after gene duplication. Instead, the change following duplication that occurred in *ScGAL1* was the acquisition of low basal expression in a medium lacking galactose and high fold expression in a medium containing galactose without loosing signalling function. Not only different species of *S.cerevisaie* lost diauxy lag [65], even different strains of *S. cerevisiae* differ in their diauxy lag [67, 68], indicating that *Saccharomyces* in general, has evolved mechanisms to fine tune its response to a medium containing glucose and galactose [48]. According to this, duplication and sub-functionalisation at the promoter level is not driven to resolve the presumptive adaptive conflict of the ancestor but to acquire additional evolutionarily advantage when *S.cerevisaie* transits from a medium rich in glucose to a medium containing galactose.

The above possibility is further strengthened by the observation that all the 12 amino acid residues of *Sc*Gal3p, which participate in binding *Sc*Gal80p are conserved in *Sc*Gal1p as well. Based on this we suggest that the reduced *Sc*Gal80p binding activity of *Sc*Gal1p as compared to *Sc*Gal3p, is dictated by other structural features of the *Sc*Gal1p, as it has to catalyse the phosphorylation of galactose as well. Thus, there is *a priori* no reason to infer that *Sc*Gal1p has been sub-functionalised partially by reducing the signalling activity. We therefore turned our focus to understand how signalling and catalytic activities are enmeshed in one single polypeptide of *Sc*Gal1p. Unlike classical allosteric enzymes [69, 70, 71], in *S.cerevisiae* galactokinase, both allosteric and orthostatic sites are one and the same. That is, galactose and ATP serve as the substrates as well as the ligands for allosteric activation.

Our strategy to address the above conundrum was to identify key residues which upon substitution would alter the protein dynamics resulting in subtle changes in the relative activities of kinase and signalling. The premise was that the protein dynamics is at play in dictating the differential functional response to the same set of molecules, that is ATP and galactose. Based on the results obtained from the analysis of *Sc*Gal1_K266R_p and *Sc*Gal1_Y441A_p, we attempt to rationalise the mechanisms of bifunctionality of *Sc*Gal1p. These two variants have improved signalling activity, but have opposite effects on the kinase function. That is, *Sc*Gal1_K266R_p gained signalling function at the cost of kinase activity, while *Sc*Gal1_Y441A_p gained both activities albeit to different extent. This clearly suggest that increase in one activity need not necessarily be at the cost of the other activity. Thus, at least in *Sc*Gal1p, there appears to be no adaptive conflict.

We suggest that in a population of active galactokinase molecules, kinase conformers and signalling conformers add up to one. That is, an increase in either kinase or signalling activity is at the cost of the other, as observed in *Sc*Gal1_K266R_ variant. If so, how do we explain an increase in both kinase and signalling activities in *Sc*Gal1_Y441A_p variant? Our analysis showed that 25.2% of *Sc*Gal1p molecules exists in open non-kinase conformation (Figure 6). It is possible that Y441A substitution results in the redistribution of open non-kinase population into closed kinase and closed non-kinase conformation but to different extent. This also explains that, Y441 *per se* is neither required for kinase activity nor for signalling. That is, if Y441 was an essential residue for the above activities, a substitution of this with Ala would have resulted in either loss or reduction in any one or both of the activities, which is not what we observe.

Above observations and the fact that Y441 is not a conserved residue points out to the following possibility. In addition to increasing the fraction of active molecules of *Sc*Gal1p as discussed above, the hydrogen bond between Y441 and Y63 restricts the frequency of transitions of *Sc*Gal1p between two functional states. In other words, Y441-Y63 H bond appears to be a bottleneck for this transition. In multiple MD simulation, we observe that during the transition of galactokinase from kinase to non-kinase or visa versa this H bond remains intact. Very rarely we observe conformations where this H bond is broken. We therefore propose, that disruption of this H bond by substituting Y441 with alanine, decreases the energy barrier and allows galactokinase to visit both the conformations with ease, resulting in an increase in both activities, in an absolute and not relative sense. Alternative explanation for the increase in both activities is that replacing a bulkier/aromatic residue (Y441) with a less bulkier/aliphatic residue (A441) could result in the phenomena described above.

The above possibility appears unlikely because of the following observations. The *Saccharomycetales* family, which is ancestral to *Saccharomycetaceae*, have either Asn or Phe instead of Tyr. Further, except, *Zygosaccharomyces bailii*, and *Lachancea fermentatti*, the rest of the species belonging to*, Saccharomycetaceae* have Tyr (Figure S12). Thus, it appears that hydrogen bond is important and not the type of substitution because most of the ancestral galactokinase sequences have Asn and not Trp or Phe. This idea is in consonance with the general concept that the ancestral proteins are functionally more plastic [72]. Second, the crystal structure of *Sc*Gal3p [38] shows that the corresponding residues form H bond. However, MD simulation shows that this H bond breaks with in few nanoseconds and remains so for the next 150 ns. According to the idea proposed here, if this H bond were to be present in *Sc*Gal3p, its signal transduction activity could have been restricted. Under similar conditions, *Sc*Gal3-SAp derivative tends to retain this hydrogen bond not as consistently as *Sc*Gal1p. Recall, that *Sc*Gal3-SAp variant has a weak galactokinase activity. Thus, it appears that the propensity for the formation of this H bond, is in some way linked to the reinstatement of kinase activity, implying that the kinase activity is established by reducing the inherent propensity of the protein to visit the signaling conformation. Based on these results, we speculate that the H bond between Y441 and Y63 is one of the important barriers for the interconversion between signalling and kinase function.

While a network of H bonds has been demonstrated to be responsible for very many functions [73, 74], hardly few studies have reported role for a H bond in the transition of protein conformation with biological implications. We came across just two reports where in a H bond is implicated in biological activity. It was reported that in Sec14p of yeast, breakage of H bond between the side chains of Q254 and D115 or the backbone H bond formed between Q254 and K116 is sufficient to bring the conformational change required to carry out the phospholipid exchange. More interestingly, this mechanism has been evolutionarily conserved in other members of the family as well [75]. A more recent study has demonstrated that a difference in the length of H bond by 0.15A, between meso and thermo ketosteroid isomerase is responsible for conferring the temperature dependence on the catalytic activity [76]. Thus, the idea that Y441-Y63 H bond could be a determinant of the conformation transition between catalysis and signalling may not be all that far fetched. If interconversion between signalling and catalysis is a biological necessity, then, that being determined by a H bond makes sense, as this mechanism provides the necessary conformational plasticity. That is, a energetically more stronger barrier would be counterproductive. A prediction from this study is that the substitution of Y441A in *Sc*Gal3p should not result in an increase in signalling. This is because, in *Sc*Gal3p, these two residues do not form a H bond in the first place, as demonstrated by our MD simulation data. However, the above data do not tell us the mechanism that triggers the interconversion, in the first place.

*Sc*Gal1_K266R_p variant differs from *Sc*Gal1_Y441A_p variant in very many ways. First, the increase in signalling activity of *Sc*Gal1_K266R_p is at the cost of kinase activity, unlike what is observed in *Sc*Gal1_Y441A_p. Second, the fold expression of *GAL1p::GFP* is clearly more than what is observed with either the *Sc*Gal1_wt_p or *Sc*Gal1_Y441A_p. Third, the steady state expression of *Sc*Gal1_K266R_p is <*Sc*Gal1_Y441A_p<*Sc*Gal1_wt_p.Fourth, the number of rapid inducers produced by *Sc*Gal1_K266R_p is more than that of *Sc*Gal1_Y441A_p. Fifth, *Sc*Gal1_K266R_p is more sensitive to galactose concentrations than either *Sc*Gal1_Y441A_p or*Sc*Gal1_wt_p. Sixth, *Sc*Gal1_K266R_p has constitutive activity. That is, it can activate the switch independent of galactose, unlike the *Sc*Gal1_Y441A_p variant. Thus, it appears that, if all other factors are held constant, an increase in signalling is at the cost of kinase and visa versa. Higher signalling activity both in terms of increase in rapid numbers as well as fold induction could be due to an increase in affinity of *Sc*Gal1_K266R_p to *Sc*Gal80p. This increase results in an increase in the half-life of the *Sc*Gal80p/*Sc*GAL1_K266R_p complex causing higher transcriptional burst size, burst frequency or both. It appears that *Sc*Gal1_K266R_p consists of conformations that can bind *Sc*Gal80p on their own (constitutive) as well as conformations that can bind *Sc*Gal80p in response to galactose and ATP. Based on the above, we speculate the K266 is a lynchpin in conferring bifunctionality to galactokinase.

The above possibility is supported by the following observation. A comparison of side chain motion of K266 (in *Sc*Gal1p), K258 (in *Sc*Gal3p) and K260 (in *Sc*Gal3-SAp) during MD simulation, suggests that this Lysine initiates/triggers the conformational transition. For example, the side chain motion of K260 of *Sc*Gal3-SAp resembles more to K266 of *Sc*Gal1_wt_p than K258 of *Sc*Gal3p. The side chain motion of K258 uncouples the catalytic residues from galactose in *Sc*Gal3p while in *Sc*Gal3-SAp it just distorts the catalytic loop. In this context, it is interesting to note that K266R substitution results in an increase in signalling at the cost of kinase activity. Arg is a conservative residue and is found in many other galactokinases such as Humans and bacteria [54], but not in fungal galactokinases.

Based on these observations, we speculate that instead of Arg at position 266, Lys is recruited to function as a swivel for conferring the ability on *Sc*Gal1p from visiting both kinase and signalling conformation (see below for more details), while H bond between Y441 and Y63 restricts this transition. As mentioned before, it is equally possible that there could be other H bonds which may serve a similar function as the H bond between Y441 and Y63. Essentially, this tug of war, that determines to what extent *Sc*Gal1p functions as a signal transducer and as a kinase. This null hypothesis is a beginning to understand the conformational transition in not only yeast galactokinase but also other moonlighting proteins where Lysine is present in the active site.

The presence of Lysine in catalytic site across protein kinases [77] seems to be a universally conserved phenomenon. A case in point is moonlighting protein kinases, which perform catalytic independent regulatory functions, consists of a lysine in their catalytic core. However, the mechanism by which they exhibit moonlighting is unclear. Interestingly it has been observed that substituting Arg for lysine residue of some protein kinase blocks catalysis, but doesn’t affect their non-catalytic functions [78, 79, 80]. It is recently reported that the moonlighting activity of ILV1 encoded threonine deaminase of *S.cerevisaie* plays a role in the regulation of gene expression in response to stress. Here, K109A substitution knocks of its catalytic activity but not its moonlighting function [81], suggesting that K109 could be playing a similar role to what we observe in galactokinase. In this context, it will be interesting to observe the features of *Sc*Gal1_K226A_p. Thus, the proposed role of Lysine is not restricted only to kinases and therefore appears to be a general feature of moonlighting enzymes containing Lysine at the catalytic core.

This study opens up a plethora of avenues to study not just conformational transition between catalysis and moonlighting activity, but also how molecular tinkering can alter the performance of *GAL* genetic switch itself. So far, the autoregulatory feed back loop of *GAL1* has been mainly studied by changing the promoter activity [82,83,84]. Now it is possible to look at these features without altering the promoter activity. In this context, it would be interesting to see whether these mutants would have any advantage in a wild type background, that is in a *GAL3+* background. For example, it has been proposed that sequestration of *Sc*Gal80p by both *Sc*Gal1p and *Sc*Gal3p give rise to co-operativity of *GAL* genetic switch [84]. Thus, if wild type *GAL1* is replaced by *Sc*Gal1_Y441A_p in a strain with wt*GAL3* allele, one would expect an increase in Hill coefficient. On the other hand, replacing*Sc*Gal1_K266R_p can have a totally different outcome because of its constitutive nature. In this case, the magnitude of the incomplete penantrance observed in the *gal3Δ* background might get exacerbated because of the wt*GAL3* mediated positive feed back loop. Alternatively, the incomplete penetrance could be completely abolished, turning on the switch completely constitutive. In either case, this phenomena would most likely make the cells less fit in an environment devoid of galactose, as the *GAL* genes would be induced with no avail. This may explain, why although Arg is present instead of Lysine in other galactokinase, in most of the fungal galactokinases Lys is preferred. At the structural level, it would be interesting to see the effect of combining K266R and Y441A (double mutant background) on signalling and catalysis. If the pathway of operation of these two variants is the same, then, there should not be any increase in signalling if these two mutations are combined. If they follow different pathway, the combined effect is expected to be more than the effect observed for the single mutations.

A fall out of the above study is our improved understanding of how trans-acting (changing activity at the level of protein) changes alone can bring about a change in the phenotypic output. That is, the incomplete penetrance exhibited by*Sc723gal3*Δ*GAL1wt* and *Sc723gal3*Δ*GAL1Y441A*is different in terms of magnitude (see table 2) but galactose dependent. However, *Sc723gal3*Δ*GAL1_K266R_* exhibits both galactose dependent and independent incomplete penetrance. In humans, incomplete penetrance underlies many diseases phenotypes and the underlying molecular mechanism are far from clear. A case in point is the incomplete penetrance exhibited by cancer stem cells [85, 86]. Cancer stem cells give rise to cancer cells at a certain frequency and this is thought to be due to secondary mutations that arise in cancer stem cells. Instead, a mechanism similar to what is described above can also be at play. That is, it is not always necessary to invoke secondary mutations to explain how cancer cells originate from cancer stem cells. According to this idea, normal cells become cancer stem cells by acquiring mutations, and start multiplying independent of the regulatory signal, similar to what we observe in the *Sc723gal3*Δ*GAL1_K266R_*. Cancer stem cells upon division, give rise to cancer cells at a certain frequency and also keep replenishing the cancer stem cells. Thus, in a way, cancer stem cells exhibit incomplete penetrance. In principle, genetic mechanism discussed in this study be operative in higher eukaryotes including humans. Thus, we generalise that a constitutive mutation in a signal transducer whose expression is otherwise driven by noise and positively autoregulated, similar to *Sc*GAL1_K266R_ variant, can give rise to a cancer stem cells which in turn give rise to cancer cells without necessarily going through secondary mutations. Such a cancer stem cell can give rise to cancer cells indefinitely in an autonomous fashion without any additional mutations. Thus, one need not have to invoke additional mutation (s) in cancer stem cells for giving rise to cancer cells and regenerating the cancer stem cells.

### Experimental methods

#### Media and growth conditions

Yeast strains grown in peptone, yeast extract and dextrose (2%) were used for transformation by the lithium acetate mediated protocol [87]. For other growth experiments, either synthetic complete medium or histidine drop out medium supplemented with glycerol 3% (vol/vol) plus lactate 2% (vol/vol) or galactose as indicated in the experiments. Ethedium bromide at a concentration of 20µg/ml was included where ever required. For growth on his d.o. medium, 3-amino-1, 2, 4-triazole to a final concentration of 10mM was used. *E.coli* were grown in LB medium and were used for propagating plasmids.

#### Strain constructions

Strains of ***S.cerevisiae*** used in this study are derivatives of *Sc723*:*MAT***a** *ade1 ile leu2-3*,*112 MEL1 ura3-52 trp1-HIII his3Δ-1lys2::GAL_UAS_-GAL1_TATA_-HIS3* [41].

Construction of <u>Sc723gal3Δ GAL1_wt_</u>

*GAL3* was deleted in Sc723 strain as described below. Sc723 was transformed with the PCR amplified Hygromycin cassette from pAG32 [88] using following primers: Forward:5’CCCCAGGTATTCATACTTCCTATTAGCGGAATCAGGAGTGCAAAAAGAGAAAATAAAAGTAAAAAGGTAGGGCAACACATAGTGCTTGCCTTGTCCCCGCCGGG3’.

Reverse:5’ACTCACCAACTTGTTTCTTTATAGAGTGTAAGAGGTATGAGTAAACTTTTAATATTTAAAGGTTGTTCCAAGAAGGTGTTTAGTGTTCGACACTGGATGGCGGCG 3’.Transformants were picked up on plates containing hygromycin at a concentration 200µg/ml. Disruption was confirmed by phenotypic and by PCR analysis using a primer upstream of *GAL3* ORF(5’AGTATTGTTTGTGCACTTGCCTGC3’.) and a primer internal to hygromycin ORF(5’CCCCGAACATCGCCTCGCTCC3’.). The resulting strain is referred as *Sc723gal3ΔGAL1_wt_*

Construction of <u>Sc723gal3Δ GAL1_K266R_andSc723gal3Δ GAL1_Y441A_</u>

*GAL1_K266R_* and *GAL1 _Y441A_*alleles were generated through SDM protocol in the plasmid pYJEBGAL1 [29], The sequences of the forward and reverse primer for generating GAL1_K266R_are:

Forward: 5’TGCGAACACCCTTGTTGTATCTAACAGGTTTGAAACCGCCCCAACCAACT3’

Reverse:5’AGTTGGTTGGGGCGGTTTCAAACCTGTTAGATACAACAAGGGTGTTCGCA3’.

The sequences of the forward and reverse primers for generating GAL1_Y441A_are Forward: 5’ATAAACTTGCCGAATGTTCTT3’

Reverse: 5’AAGAACATTCGGCAAGTTTAT3’

Mutations were confirmed by sequencing from First Base (Malaysia). The site directed mutant alleles available in the form of plasmids, as described above, were then transferred to the *GAL1* genomic locus of a *Sc723gal3ΔGAL1*using *delete perfecto* method [89] as described. A CORE cassette (CORE cassette consists of selectable/reporter marker, i.e. KanMx4 which provides resistance against G418) and a counter selectable/selectable marker, i.e. KlURA3 which allows auxotrophic ura3 strain to grow on ura d.o. medium and be counter selected on 5-FOA.) was integrated into *Sc723gal3Δ*. The CORE cassette was inserted either at the point of desired substitution or at a position close to the desired substitution.

Following forward and reverse CORE primers were used for generating the recipient for introducing the *GAL1_K266R_* mutation in to the genome.

Forward:5’TACTCCGTTTAAATTTCCGCAATTAAAAAACCATGAAATTAGCTTTGTTATTGCGAACACCCTTGTTGTATGAGCTCGTTTTCGACACTGG3’.

Reverse:5’AACATTTGCAGCTGTAGTGACTTCTACCACTCTTAAATTATAGTTGGTTGGGGCGGTTTCAAACTTGTTAGTCCTTACCATTAAGTTGATC3’.

Following forward and reverse CORE primers were used for generating the recipient for introducing the *GAL1_Y441A_* mutation in to the genome.

Forward:5’ATTCACAAGAGACTACTTAACAACATCTCCAGTGAGATTTCAAGTCTTAAAGCTATATCAGAGGGCTAAGCGAGCTCGTTTTCGACACTGG3’.

Reverse:5’GTCGGCAGTAAAGCTCGCTGTAGTCATTAATTTCACAGCCTTCAAGACTCTTAAAGATTCAGAATACACATTCCTTACCATTAAGTTGATC3’.

These CORE containing strains were transformed with the PCR amplified mutated *GAL1_K266R_* and *GAL1_Y441A_* so that *GAL::CORE* ORF gets replaced with mutated *GAL1* ORF. Cells which undergo this replacement are isolated by negative selection of 5-FOA media and sensitivity on G418 containing media.

The PCR amplification of the mutated *GAL1_K266R_* and *GAL1_Y441A_* ORF was done using following primers.

Forward: 5’ATGACTAAATCTCATTCAGA3’

Reverse: 5’TTATAATTCATATAGACAGCT3’

The introduction of the desired mutation in *GAL1* was ascertained by sequencing the *GAL1* locus from the genomic DNA isolated from the putative *Sc723gal3ΔGAL1_K266R_* and *Sc723gal3ΔGAL1_Y441A_*.

Following primers were used for sequencing.

Forward: 5’TTATTTCTGGGGTAATTAAT3’

Forward:5’ATGACTAAATCTCATTCAGA3’

Forward: 5’TCTGTGAGGGTGATGTACCAA3’

Forward:5’ATTTAAGAGTGGTAGAAG3’

Forward:5’TGGTTGTACTGTTCACTTGGTTC3’

Reverse: 5’TTATAATTCATATAGACAGCT3’

Construction ofSc723gal3Δ GAL1_wt_ gal80::LEU2, Sc723gal3Δ GAL1_K266R_ gal80::LEU2 and Sc723gal3Δ GAL1_wt_ gal80::LEU2

GAL80 was disrupted inSc723gal3ΔGAL1_wt_, Sc723gal3ΔGAL1_K266R_ and

*Sc723gal3*Δ*GAL1_Y441A_*. by integrating *gal80:LEU2* cassette. The *GAL80* deletion cassette was PCR amplified from genomic DNA of BY2685 [67] using the following primer sequences (Forward: 5’CCTCCTCCAGATGGAATCCCTTCCATAG3’ Reverse:

5’GCAAACCTATCACCCGGTGATAACAGC3’.

*GAL80* disruption in these strains was confirmed by testing the sensitivity towards 2 deoxygalactose as well as by PCR amplification using the external primer upstream of *GAL80* ORF and primer internal to *LEU2* ORF.

The sequences of external and internal primer are as follows:

External: 5’ GCCTGTCTACAGGATAAAGAC3’

Internal: 5’CGCCAAGCTTGATATGAG3’

Construction of *Sc723gal3Δ GAL1_wt_ GAL1p:GFP_;_Sc723gal3Δ GAL1_K266R_ GAL1p:GFPand Sc723gal3Δ GAL1_Y441A_ GAL1p:GFP* Sc723 with the integrated *GAL1p::GFP* cassette was already available in the laboratory. *GAL1p::GFP* reporter cassette was introduced by transforming *Sc723gal3*Δ*GAL1_wt_*, *Sc723gal3*Δ*GAL1_K266R_* and *Sc723gal3*Δ*GAL1_Y441A_* to uracil prototrophy withYIp*lac*211*GAL1::GFP* [57] after digesting with *Eco*RV.

Protocol for rapid inducer assay using growth on galactose as well as growth on histidine drop out media was carried out as described in [57]. FACS analysis was carried out as described [49] except the model BD FACS ARIA-1was used. Result of a typical experiment is presented and is an average of triplicate samples. FACS analysis was repeated three times. Galactokinase assay was carried out as described [27]. Enzyme activity was measured in the linear range of enzyme concentration. Each data point is an average of three independent experiments and each sample was in triplicate. For western blot polyclonal antibodies raised against *Sc*Gal3p was used at a dilution of 1:7500 as described [47]. Experiments were repeated multiple times at different protein concentrations. Band intensity was measured using the ImageJ software.

### Computational methods

#### Conformational Sampling

Multiple MD simulation trajectories were seeded starting from diverse conformers of ScGal1p. These diverse conformers were generated by a hybrid method of anisotropic network model combined with Generalized born implicit solvent based MDsimulations [90]. The conformers generated using this approach were clustered together based on their mutual RMSD values and from every cluster a representative structure was built. From these representative structures, only those conformers that differed from other represenative structures atleast by an RMSD of 2Å, were selected. These conformers were used to further seed long MD simulation trajectories.

#### Molecular Dynamics Simulations

MD simulations of ScGal1p(i.e. ScGal1p bound to ATP and Galactose)was run in NAMD package at a temperature of 300K with periodic boundary conditions using CHARRM force field and Tip3p water model. The solvent box size was chosen such that the distance between protein to the wall was 30A^0^. Covalent bonds involving Hydrogen atom were made rigid using Shake algorithm. The cut-off for non-bonded interactions was set up at 12A^0^and electrostatic interactions were modeled using PME method. The simulation was done at 1bar in an NPT ensemble using a constant temperature of 300K. The scripts for analysis of trajectories were generated in-house using MATLAB, python and VMD.

## Acknowledgements

Financial assistance is provided by the Department of Science and technology through a grant (EMR/2016/005027) awarded to Jayadeva Bhat. NandineeGiri is a recipient of CSIR, Govt of India SRF fellowship. We thank computational facility provided by CDAC, Pune. FACS facility of the dept was funded by FIST program of Dept of Science and Technology..We thank Prof. RanjithPadinhateri for providing computational facility. We also thank, Prof. Punekar, Prof. Balaji and Prof. Bhaumik, Prof. Hossur for discussions and critical comments.We thank Prof. Saini for critically reading the manuscript.

