## Supplementary information for "Yeast galactokinase in closed conformation can switch between catalytic and signal transducer states"

**Supplementary information:**
**Statistical coupling analysis.**
The algorithm of SCA was adopted from Lockless & Ranganathan, 2009. Independent sectors were obtained by the spectral decomposition of Positional correlation matrix. The positional correlation matrix was constructed using the Kullback-Leibler relative entropy (KLE). The dataset of 82 galalctokinase like sequences belonging to *Saccharomycotina* group was used for this purpose.

**SCA analysis:**

1. **Construction of Multiple sequence alignment:**
   Dataset of 82 galactokinase like sequences belonging to *Saccharomycotina* group was used. Multiple sequence alignment of these sequences was carried out. *Sc*Gal1p sequence was taken as the reference sequence. That is, the numbering of the positions of the alignment was done as per the amino acid positions of *Sc*Gal1p sequence. The multiple sequence alignment was truncated to include only those positions whose gap frequency was lesser than 20%. This truncation step resulted in inclusion of only 497 positions from 528 positions of *Sc*Gal1p sequence. This means that there were 31 positions in the alignment where out of 82 sequences, gaps were present in more than 16 sequences. The truncation of the alignment to exclude highly gapped positions of the alignment is a step recommended by the authors of the SCA technique. This is to ensure that the calculations are done majorly at the non-gapped positions of the alignment. This multiple sequence alignment of 82 sequences and 497 positions is represented by a binary array $x_{is}^{(a)}$ where $x_{is}^{(a)} = 1$ if sequence 's' has amino acid 'a' at position 'i' and 0 otherwise. The size of this binary array is 82 rows X 9940 columns (i.e.82 sequences(rows) where each amino acid position is represented by 20 amino acids, which leads to 20*497 = 9940 columns). The sequence similarity matrix is calculated by the following equation.
   S = msa20*msa20'/N_pos

   Where S = Sequence similarity matrix

   msa20 = Binary array with 82 rows and 9940 columns(see above text).

   msa20' = Transpose of the binary array

   N_pos = Number of amino acid positions in the alignment. In this case it is 497.
   The sequence similarity histogram is shown in figure S1A.

2. **Positional correlation matrix:**

   The conservation score of each amino acid position was measured by Kullback-Leibler relative entropy (KLE) which is calculated as follows:

   $$D_i = \sum_{(a=0:20)} f_i^{a} * \ln(f_i^{a}/q^{a})$$

   Where $f_i^a$ = frequency of amino acid 'a' at position 'i'

$f_i^0$ = frequency of gap at position 'i' and is given by $f_i^0 = 1 - \sum_{(a=1:20)} f_i^a$

$q^a$= Background probability of amino acid 'a' which is calculated from the mean frequency of amino acid 'a' in non-redundant protein database. This background probability score is provided with the SCA toolbox.

$q^0$ = Background probability of gaps.

The positional conservation score profile is shown in figure S1B. Based on the above positional conservation score of each amino acid position, a positional correlation matrix was constructed.

3. **Spectral decomposition of positional conservation matrix.**
The magnitude of the eigen vectors obtained from the diagonalization of this matrix form the basis of sector identification. In order to ensure that the eigen vectors taken into consideration could identify functionally relevant sectors, the sequence alignment was randomized and a positional correlation matrix from the randomized alignment was constructed. The randomization process was done at least 100 times. This process resulted in identification of those eigen values of the positional correlation matrix of the real sequence alignment which were not sampled in the randomized alignments. (Figure S1, panel C) This process resulted in identification of 7 significant eigen vectors from which 7 over-lapping sectors were identified. To ensure that these sectors were independent of each other i.e. there would be no linear relationship between the sectors (this means that each sector would serve as a separate functional unit of ScGal1p), these eigen vectors were subjected to independent component analysis which identified 7 non-overlapping statistically significant independent sectors (Figure S1,panel D).

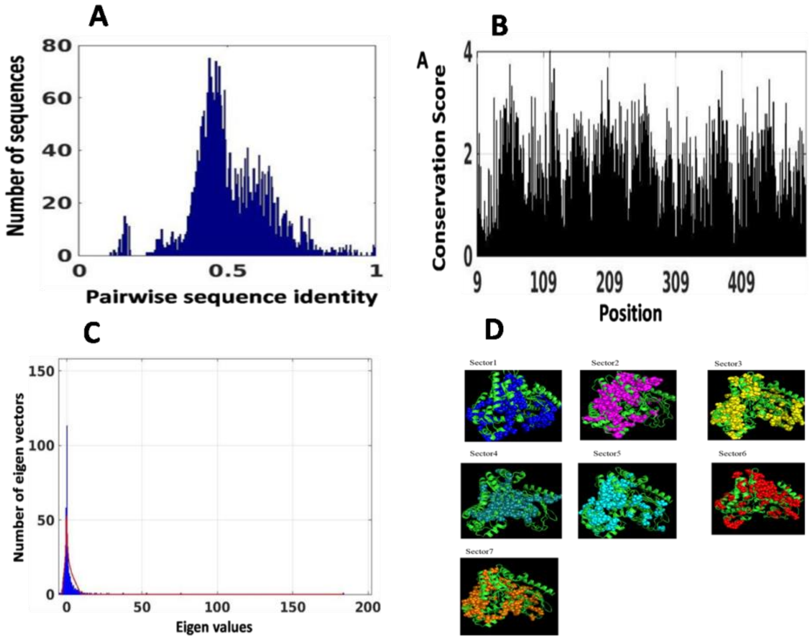


**FigureS1: SCA analysis for galactokinase sequences belonging to *Saccharomycotina* group.** Panel A shows the distribution of pairwise sequence similarity of galactokinase like sequences belonging to *Saccharomycotina* group (the list of sequences used for this analysis is available on request). Panel B shows the positional conservation score of amino acid positions of *Sc*Gal1p. Panel C shows the distribution of eigen values of positional correlation matrix. Red curve indicates the distribution of eigen values of the positional correlation matrix of original alignment. The blue shaded region represents the distribution of eigen values of the positional correlation matrix of the randomized alignments. Panel D shows the seven sectors.

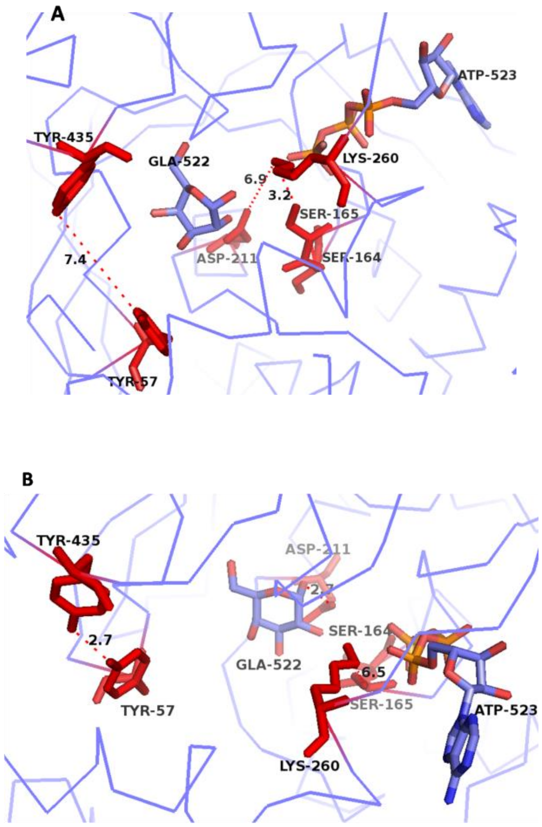

**FigureS2:Kinase and Non-kinase states of *Sc*Gal3p-SA during 150ns simulation.** Panel A shows the structural orientations of Y57, Y435, K260, S165, D211,galactose and ATP in initial structure of *Sc*Gal3p-SAp. Panel B shows the structural orientations of Y57, Y435, K260, S165, D211, galactose and ATP in*Sc*Gal3p-SAp during the later stages of 150ns simulation(i.e when distance b/w anomeric hydroxyl group of galactose and D211 reduces to less than 3.5A).

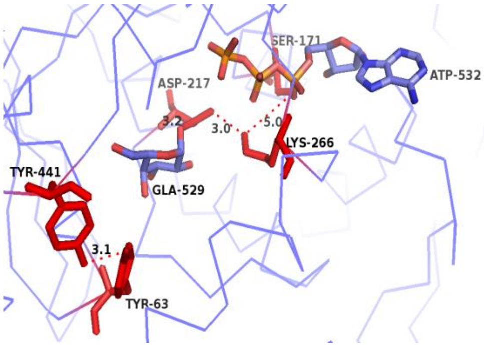


**Figure S3: Conformation of *Sc*Gal1p during 150ns simulation.** This panel shows the structural orientations of Y63, Y441, K266, S171, D217,galactose and ATP in *Sc*Gal1p during 150ns simulation.

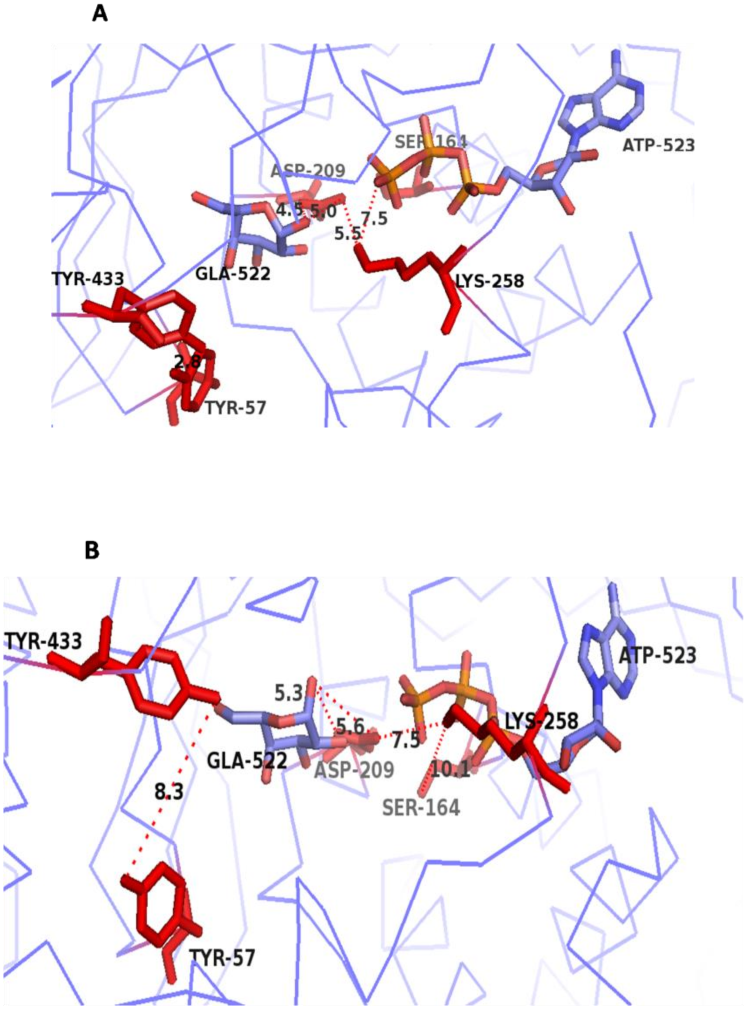


**Figure S4:Conformation of *Sc*Gal3p during 150ns simulation.** Panel A shows the structural orientations of Y57, Y433, K258, S164, D209,galactose and ATP in crystal structure of *Sc*Gal3p. Panel B shows the structural orientations of Y57, Y433, K258, S164, D209, galactose and ATP in *Sc*Gal3p as the simulation progresses.

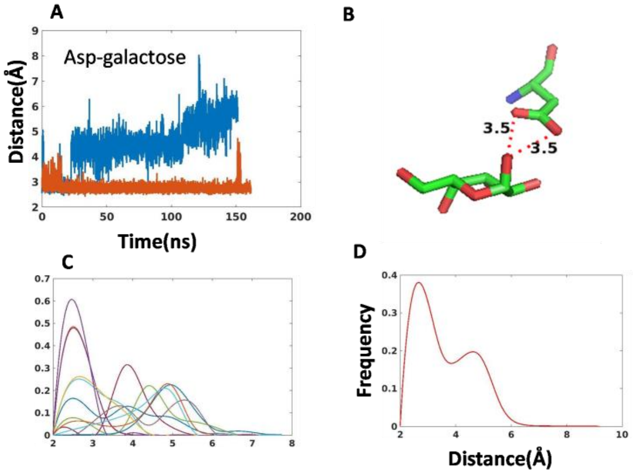


**Figure S5:*Sc*Gal1p samples kinase and non-kinase states based on the distance between D217 and anomeric hydroxyl group of galactose as order parameter.** Panel A shows the distance between catalytic aspartate residue (D217) and anomeric hydroxyl group of galactose plotted as a function of time. Orange and blue refers to the trajectories starting from two different seed conformations. Panel B shows the distance between anomeric hydroxyl group of galactose and D217 in crystal structure of *Sc*Gal1p. Panel C shows the distribution of distance between anomeric hydroxyl group of galactose and D217 for each of the 19 trajectories separately. Panel D shows the distribution for all the trajectories clubbed together.

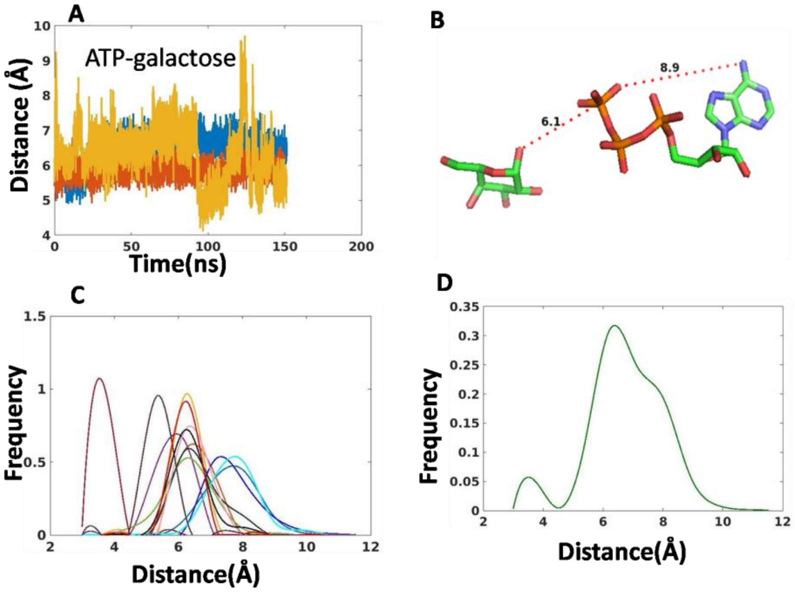


**Figure S6:*Sc*Gal1p samples kinase and non-kinase states based on the distance between terminal phosphate of ATP and anomeric hydroxyl group of galactose .**Panel A shows the distance between terminal phosphate of ATP and anomeric hydroxyl group of galactose plotted as a function of time.Orange, blue and yellow refers to thetrajectories starting from three different seed conformations.Panel B shows the distance between anomeric -OH group of galactose and terminal phosphate of ATP in crystal structure of *Sc*Gal1p. Panel C shows the distribution of distance between anomeric -OH group of galactose and terminal phosphate of ATP for each of the 19 trajectories separately. Panel D shows the distribution for all the trajectories clubbed together.

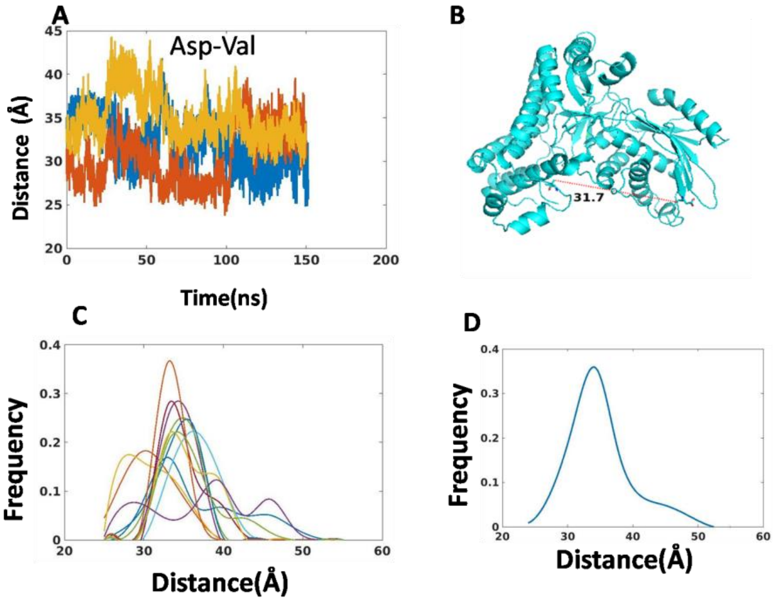


**Figure S7:*Sc*Gal1p samples open and closed states.** Panel A shows the distance between residues V383 and D106 plotted as a function of time. Orange, blue and yellow refers to thetrajectories starting from three different seed conformations.Panel B shows the distance between V383 and D106 in crystal structure of *Sc*Gal1p. Panel C shows the distribution of distance between for each of the 19 trajectories separately. Panel D shows the distribution for all the trajectories clubbed together

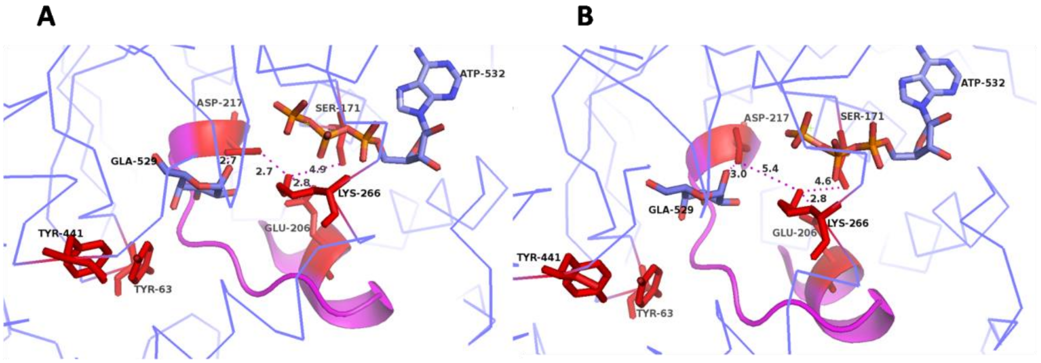


**Figure S8:Kinase states of *Sc*Gal1p.** Panel A corresponds to the average structure of *Sc*Gal1p when the distance between K266 and D217 is less than the 4A. Panel B corresponds to the average structure of *Sc*Gal1p when the distance b/w K266 and D217 is more than the 4A. The functional groups analysed in this study is highlighted in red and megenta.

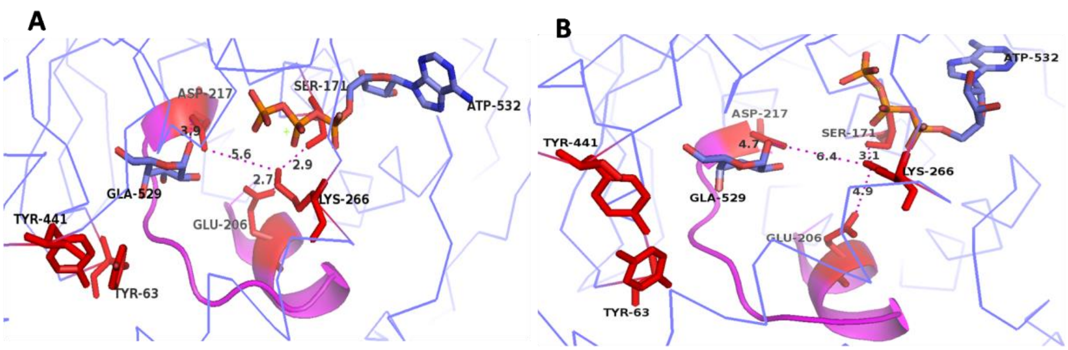

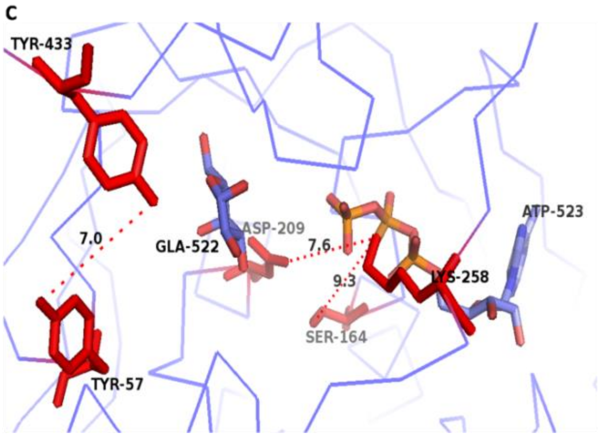


**Figure S9: Signal transducer state of *Sc*Gal1p and *Sc*Gal3p.** Panel A corresponds to the average structure of *Sc*Gal1p when the distance between K266 and D217 is less than the 6A. Panel B corresponds to the average structure of *Sc*Gal1p when the distance betweem K266 and D217 is more than the 6A. Panel C corresponds to the structure(snapshot) of *Sc*Gal3p observed during later stage 150ns simulation when there is a decoupling of galactose from catalytic loop. The functional groups analysed in this study is highlighted in red and magenta.

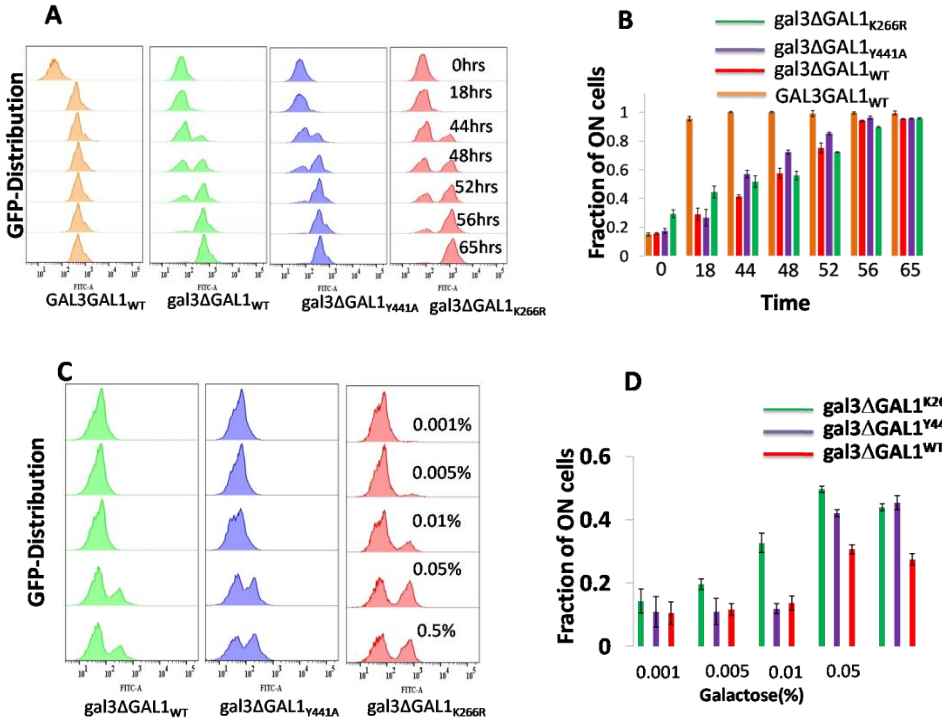


**FigureS10: Distribution of GFP expression in indicated starins.** Panel A shows the thedistribution of *GAL1*::GFP expression in cells of the indicated strains as a function of timeafter exposing the cells to 0.5% galactose. Panel B shows fraction of ON cells as a function of time.Panel Cshows thedistribution of *GAL1*::GFP expression in cells of the indicated strains as a function of different galactose concentration. Panel D shows the fraction of ON cells as a function of concentration of galactose.

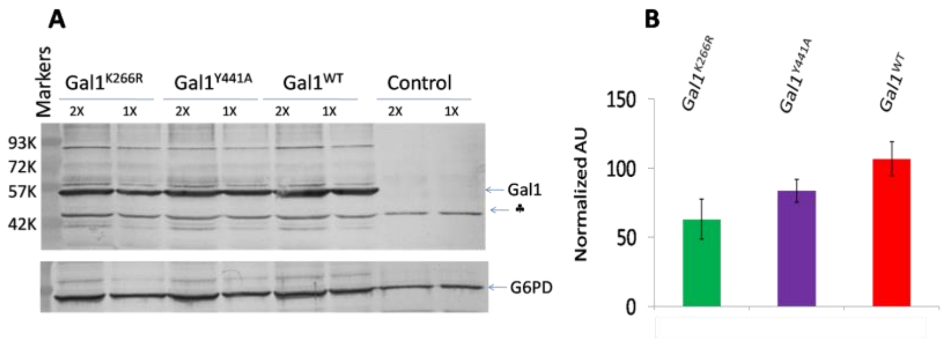


**Figure S11:Western blot analysis of wild type and the mutant GAL1 proteins.** Panel Arepresents the extract obtained form a wild type strain grown in glycerol lactate. ♣Indicates a nonspecific band picked up by the antibody against galactokinase. Lower panel represents the loading control. 'X' represents the fold dilution of the extract loaded. Right panel represents the intensity in AU normalized to 100% of the loading control.

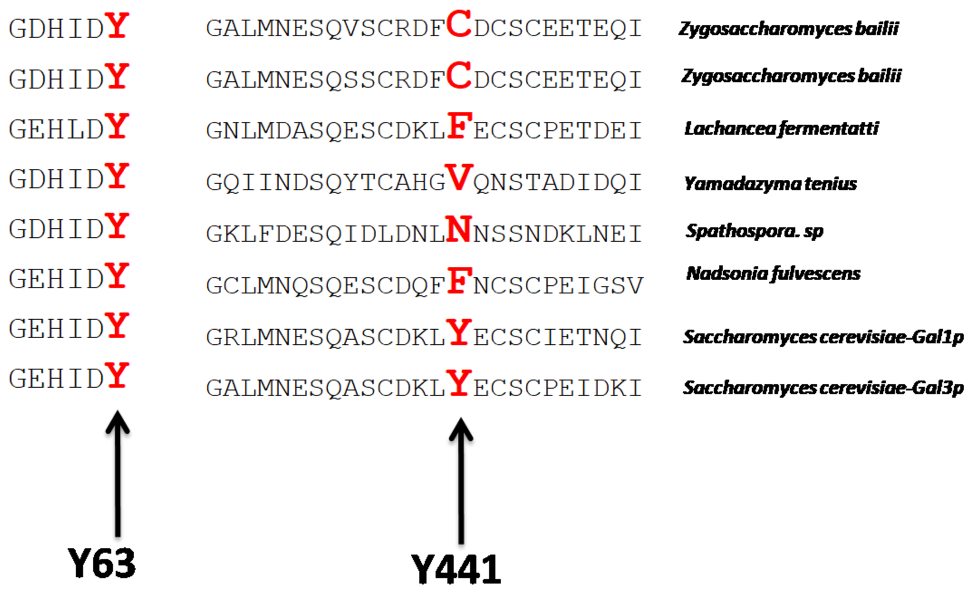


**Figure S12: Multiple Sequence alignment of the indicated species showing the sequence corresponding to *Sc*Gal1p-Y441 and *Sc*Gal1p-Y63**

**Table S1: Distribution of mutationally identified residues into different sectors.**

| Sector | Number of substitutions |
|---|---|
| 1 | 2 |
| 2 | 2 |
| 3 | 1 |
| 4 | 4 |
| 5 | 6 |
| 6 | 3 |
| 7 | 1 |

**Table S2: List of amino acid residues present in different sectors**

| Sector | Residues |
|---|---|
| Sector1 | 40,90,92,93,103,105,106,118,121,122,124,157,176,205,231,232239,241,258,260,262,272,273,282,296,302,308,345,399 411,437,441,443,445,447,465,473,474,489 |
| Sector2 | 356,382,383,388,393,395,396,397,403,404,427,433,444,452,454,459,462,464,466,467,468,469,471,472,493,510,517,519 522 |
| Sector3 | 22,44,56,65,89,102,107,110,111,119,139,143,150,186,209,248,281,286,313,317,318,321,327,342,359,368,369,386,389 407,409,419,478,520, |
| Sector4 | 50,83,117,125,227,229,256,263,264,288,294,346,385,390,394,398,429,430,451,470,494 |
| Sector5 | 9,36,49,53,55,60,61,62,63,66,67,68,69,70,95,100,116,126,127,128,129,169,173,174,210,211,214,216,218,221,230,234, 235,238,243,266,269,271,274,275,277,278,280,284 |
| Sector6 | 43,45,64,71,75,76,77,88,104,113,155,158,160,163,166,171,172,187,194,200,220,261,279,304,312,337,341,354,366,371, 420,491,497,505,506,509,518,526 |
| Sector7 | 11,12,14,46,73,81,164,185,195,201,202,203,204,207,208,225,246,268,311,315,331,332,336,362,405,461 |
